# Long-lived central memory γδ T cells confer protection against murine cytomegalovirus reinfection

**DOI:** 10.1101/2022.08.05.502906

**Authors:** Nathalie Yared, Maria Papadopoulou, Sonia Netzer, Laure Burguet, Lea Mora Charrot, Benoit Rousseau, Atika Zouine, Xavier Gauthereau, David Vermijlen, Julie Déchanet-Merville, Myriam Capone

**Affiliations:** Bordeaux University, CNRS, ImmunoConcEpt, UMR 5164, 33076 Bordeaux, France.; Department of Pharmacotherapy and Pharmaceutics and Institute for Medical Immunology, ULB Center for Research in Immunology (U-CRI), Université Libre de Bruxelles (ULB), Belgium.; Bordeaux University, Service Commun des Animaleries, 33076 Bordeaux, France; Bordeaux University, CNRS UAR3427, INSERM US05, FACSility, TBM Core, 33076 Bordeaux, France.; Bordeaux University, CNRS UAR3427, INSERM US05, OneCell, RT-PCR and Single Cell Libraries, TBM Core, 33076 Bordeaux, France.

## Abstract

The involvement of γδ TCR-bearing lymphocytes in immunological memory has gained increasing interest due to their functional duality between adaptive and innate immunity. γδ T effector memory (TEM) and central memory (TCM) subsets have been identified, but their respective roles in memory responses are poorly understood. In the present study, we used subsequent mouse cytomegalovirus (MCMV) infections of αβ T cell deficient mice in order to analyze the memory potential of γδ T cells. As for CMV-specific αβ T cells, MCMV induced the accumulation of cytolytic, KLRG1+CX3CR1+ γδ TEM that principally localized in infected organ vasculature. Typifying T cell memory, γδ T cell expansion/proliferation in organs and blood was higher and more efficient after secondary viral challenge than after primary infection. Viral control upon MCMV reinfection involved the T-cell receptor, and was associated with a preferential amplification of private and unfocused TCR δ chain repertoire as evidenced by next generation sequencing. The γδ T cell secondary response to MCMV was composed by a combination of clonotypes expanded post-primary infection and, more unexpectedly, of novel expanded clonotypes. Finally, Long-term-primed γδ TCM cells, but not γδ TEM cells, protected T cell-deficient hosts against MCMV-induced death upon adoptive transfer, probably through their ability to survive and to generate TEM in the recipient host. Overall, our study uncovered memory properties of long-lived TCM γδ T cells that confer protection in a chronic infection, highlighting the interest of this T cell subset in vaccination approaches.

**AUTHOR SUMMARY:** Cytomegalovirus (CMV) is a widespread, latent virus that can cause severe organ disease in immune-compromised patients. Anti-CMV memory immune responses are essential to control viral reactivation and/or reinfection events that commonly take place in solid organ transplantation. The role of γδ T-cell receptor bearing lymphocytes could be crucial in this context where immunosuppressive/ablative treatments cause suboptimal and/or delayed αβ T cell responses. Here we asked whether γδ T cells could compensate for the absence of αβ T cells in the long-term control of mouse CMV infection. Three months post-primary viral challenge in αβ-T cell deficient mice, γδ T cells displayed similar features as cytolytic, CMV-specific αβ CD8 T cells. We showed that previous priming with CMV endowed γδ T cells with an enhanced antiviral potential and that long-term maintenance of γδ-mediated antiviral protection was dependent on γδ central memory T cells (TCM). The γδ T cell response to a secondary CMV challenge was dependent on γδ TCR-signaling and generated a private TCR δ repertoire as observed in human. Our results sustain the adaptive-like properties of these unconventional T cells and reveal the interest of targeting γδ TCM subset in novel antiviral vaccination approaches.

## INTRODUCTION

The concept of immunological memory has been challenged in recent years. It was described as an acquired, multidimensional and evolutionary process that can no longer be solely related to vertebrates and adaptive immunity. With regards to the host capacity to improve survival upon secondary infection, immunological memory implicates both the innate and adaptive arm of the immune system (reviewed in [1–3]).

Prototypical adaptive memory cells are CD8+ αβ T lymphocytes, composed of various populations harboring distinct migratory, proliferative and effector properties: CCR7+CD62L+CD27+ Central Memory T cells (TCM), mostly found in secondary lymphoid organs, and CCR7-CD62L-CD27-Effector Memory T cells (TEM), among which CX3CR1+ circulating TEM and CX3CR1-CD103+CD49a+CD69+ Tissue-Resident Memory T cells (TRM). In the case of resolving infections, CD8 TEM progressively decline unless submitted to antigenic re-challenge. When the pathogen persists, these cells are retained and acquire specific features reminiscent to the nature of infection. In chronic infections (LCMV clone 13 in mice or HIV, HCV in humans), antiviral CD8 TEM display an increased expression of inhibitory receptors such as PD-1. In latent infections however (herpesviruses), such “exhausted” phenotype is uncommonly found, at least among circulating CD8 TEM (For reviews see [4–7]).

Among latent β-herpesviruses, cytomegaloviruses (CMVs) have drawn great attention due to the deleterious effects of CMV infection in immune-compromised individuals. In healthy subjects, remodeling of the CD8+ αβ TCR repertoire occurs over time during latency, with an increased frequency of cells specific for a few viral epitopes, a phenomenon referred to as memory inflation. Long-term CMV-induced CD8 T cells express a TEM KLRG1+ phenotype. In humans, they use the longer CD45 isoform (CD45RA), reminiscent of highly differentiated cells. Inflationary CD8+ αβ T cells circulate in the blood, home to peripheral tissues and show increased expression of cytolytic proteins. They are assumed to maintain robust function and prevent viral spread upon reactivation (for review see [8–11]). The crucial role of long-lasting memory to human CMV (HCMV) is exemplified in solid organ transplantation (SOT), where seronegative recipients (R-) receiving a CMV+ graft (D+) are at higher risk of developing CMV disease than seropositive recipients (R+).

Another population expressing somatically diversified, T-cell antigen (Ag) receptors (TCR) is delineated by MHC-unrestricted, γδ TCR-bearing lymphocytes, which recognize ubiquitous stress-induced self ligands [12]. Their implication in immunological memory has been put forward in recent years, in different settings including pathogen infection and autoimmunity (reviewed [13–15]). Because of their dual nature, γδ TCR-mediated memory responses display both innate-like and adaptive-like characteristics [16–18]. In fact, γδ T cells are composed of so-called innate-like subsets with invariant and public (shared between individuals) TCR repertoires, as well as adaptive-like subsets with unfocused and highly private TCR repertoires. Innate-like γδ T lymphocytes are preprogrammed before birth and thus display rapid effector function in periphery. On the other hand, adaptive-like γδ T cells (which comprise mouse Vγ1+ and Vγ4+ and human non-Vγ9Vδ2 T cell subsets) exit the thymus after birth with a naïve phenotype. Reviewed in [19–21].

Some years ago, our group identified HCMV as a major driver of non-Vγ9Vδ2 clonal expansions in peripheral blood of renal transplant recipients, that was further on confirmed by other teams in different settings including allo-hematopoietic stem cell transplantation (HSCT) [22] (reviewed in [23, 24]). A γδ T cell contribution to the anti-viral response was suggested by the concomitant diminution of viremia [25, 26]. The protective role of γδ T cells was evidenced thanks to the mouse model. Concomitantly to the group of Winkler, we showed that γδ T cells can compensate for the absence of αβ T cells and confer protection to mouse CMV (MCMV) (in TCRα^-/-^ or CD4-depleted CD8^-/-^JHT mice)[27, 28]. In TCRα^-/-^ mice, the acute phase response to MCMV engaged Vγ1+ and Vγ4+ T cells that acquired a CD44+CD62L-(TEM) phenotype alongside infection resolution. To which extent and for how long previous contact with CMV endows γδ T cells with an increased antiviral reactivity remains to be clarified.

As observed for HCMV-specific inflationary CD8+ αβ T lymphocytes, long-term HCMV-induced non-Vγ9Vδ2 T cells express a TEMRA (CD45RA+CD27-CD62L-CCR7-) KLRG1+ phenotype. Moreover, transplant patients experiencing a secondary infection (D-R+, *i.e.* HCMV-seropositive recipients receiving graft from a HCMV-seronegative donor) show a faster recall response of TEMRA γδ T cells and better infection resolution comparatively to patients experiencing a primary infection (D+R-) [26, 29]. These results strongly suggest that γδ T cells have a memory potential against CMV. In this study, we addressed this issue using our previously set up TCRα^-/-^ mouse model that allows experimental re-infection and adoptive transfer experiments. Our results show the preferential amplification of private TRD (TCR δ chain) repertoire upon MCMV reinfection, and highlight the crucial role of CD44+CD62L-γδ TCM for long-term maintenance of γδ-mediated antiviral protection.

## RESULTS

### 1. γδ T cell response to secondary MCMV challenge is faster, higher and more efficient than during primary infection

To test whether γδ T cells have a memory potential against MCMV, we used subsequent infections of αβ T cell deficient mice, which allowed direct demonstration of the protective antiviral function of γδ T cells [27]. Three months after a first contact with MCMV (*i.e.* time delay required for latency in wild type mice), TCRα^-/-^ mice were re-challenged with similar doses of MCMV, and the secondary γδ T cell response was analyzed in the liver and lung. Age-matched control mice were primarily infected concomitantly (Fig 1A). When looking at γδ T cell numbers, a more rapid and higher increase of γδ T lymphocytes was observed after re-infection when compared to primary infection (Fig 1B, upper panels). The highest count was obtained at day 1 (d1) for liver and d7 for lung in re-infected mice, comparatively to d7 for liver and d14 for lung during primary infection. Concerning viral loads, DNA copies of MCMV were still detected in organs from d92-infected TCRα^-/-^ mice (Fig 1B, lower panels). However, after re-infection, they remained stable and even slightly decreased by contrast to the increase observed during primary infection.

**Fig 1.**
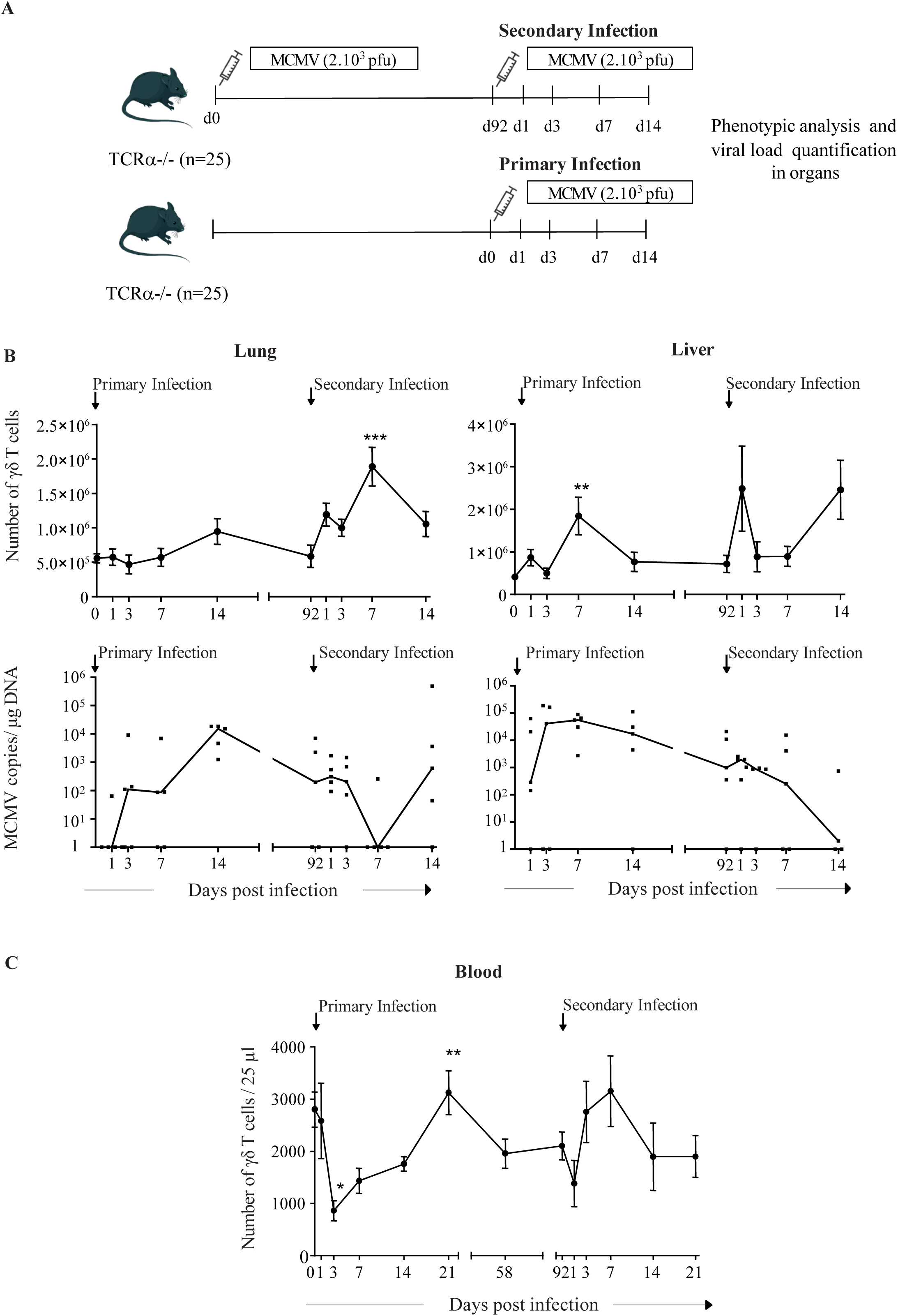
γδ T cell response to secondary MCMV challenge. (A) Experimental scheme: analyses in organs. TCRα-/- mice (n=25) were infected with MCMV (2.10^3^ PFU) at day 0, or left uninfected (n=25). 3 months later, uninfected mice were primarily infected, and infected mice were re-challenged with similar dose of MCMV. The kinetic of the γδ T cell response was analyzed in the liver and lung and depicted longitudinally although mice (n=5) were sacrificed at indicated time points. (B, upper panels) Total number of γδ T lymphocytes in organs. Data represent the mean +/- SEM of cell counts from 5 mice (1-way ANOVA). (B, lower panels) Viral loads/ 1 µg of total DNA for individual mouse that were quantified in organs by qPCR. The median is shown by a line. (C) Longitudinal analysis of γδ T cells in blood. TCRα-/- mice (n=10) were infected with MCMV (2.10^3^ PFU) at day 0, then re-challenged at day 92 with similar dose of MCMV. Mice were bled at the indicated time points. Mean absolute numbers +/- SEM of γδ T lymphocytes in 25 µl of blood are shown. Friedman statistical test was used with d3 (post-primary) and d1 (post-secondary infection) as references. (A-C) The experiment was repeated twice with comparable results.

In order to converge to analyses done in humans, we monitored the kinetic of γδ T lymphocytes longitudinally, in blood samples from TCRα^-/-^ mice taken post primary and secondary MCMV-infection. In accord with previous findings, the first phase of infection was marked by a drop in γδ T cell numbers, likely related to virus-driven lymphopenia (Fig 1C) [28, 30]). Then, γδ T cell counts increased back to normal levels, reaching a peak at day 21. Interestingly, maximal numbers of γδ T cells were obtained at day 7 post-reinfection, i.e. much earlier than after primary MCMV encounter.

Collectively, our data depict a more rapid and more efficient γδ T cell response to MCMV after secondary challenge compared to the primary response, which are typical of T cell memory.

### 2. KLRG1+ γδ TEM dominate the γδ T cell secondary response to MCMV

To characterize the memory γδ T cell subsets involved in the recall response to MCMV, we followed the kinetic of γδ T naïve (CD44-), TCM (CD44+CD62L+) and TEM (CD44+CD62L-) along the course of the experiment depicted in Fig 1A. Due to the broad antigen specificity of γδ T cells and the presence of innate-like γδ T cells in organs [31], an important proportion of γδ TEM but few naïve γδ T cells were evidenced in control uninfected mice (d0, S1A Fig). From d0 to d92, γδ TEM percentages increased concomitantly to a decrease of TCM. Notably (Fig 2A), γδ TEM numbers tripled up after secondary infection in organs (d7 for lung and d1 for liver), and blood (d7). Thus, the kinetic response of TEM in organs and blood (Fig 2A) followed that of total γδ T cells (Fig 1B, upper panels and Fig 1C), once again typifying T cell memory.

**Fig 2.**
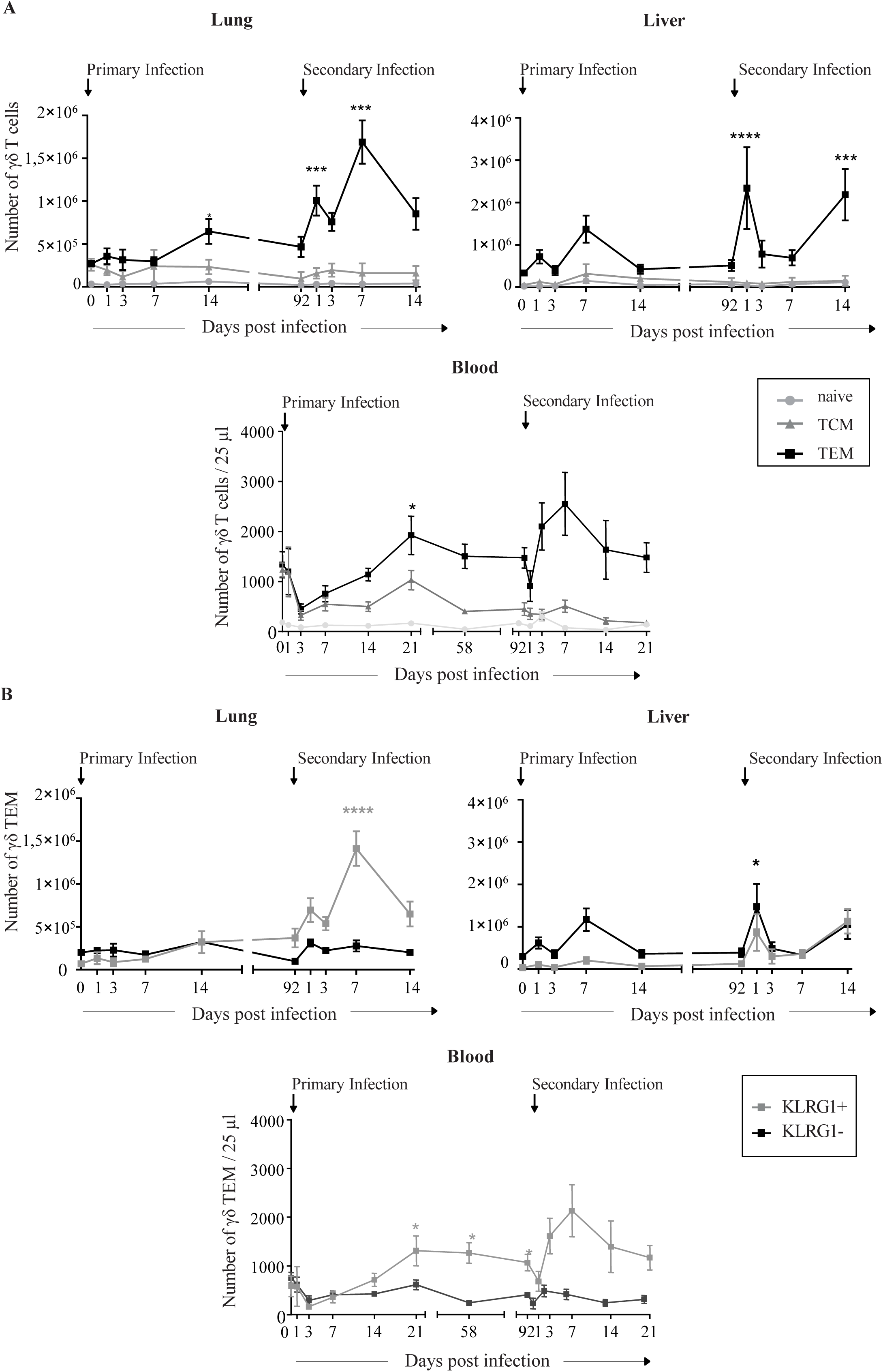
KLRG1+ γδ TEM dominate the γδ T cell secondary response to MCMV. (A) Kinetic evolution of naïve (CD44-CD62L+), TEM (CD44+CD62L-) and TCM (CD44+CD62L+) γδ T cell subsets. (A, upper panel) analysis in organs. TCRα-/- mice (n=25) were infected with MCMV (2.10^3^ PFU) at day 0, or left uninfected (n=25). 3 months later, uninfected mice were primarily infected, and infected mice were re-challenged with similar dose of MCMV. The kinetic of the γδ T cell response was analyzed in the liver and lung and depicted longitudinally although mice (n=5) were sacrificed at indicated time points. Data represent the mean +/- SEM of cell counts from 5 mice. (1-way ANOVA) (A, lower panels) Analysis in blood. TCRα-/- mice (n=10) were infected with MCMV (2.10^3^ PFU) at day 0, then re-challenged at day 92 with similar dose of MCMV. Mice were bled at the indicated time points. Mean absolute numbers +/- SEM of γδ T lymphocytes in 25 µl of blood are shown. Statistical test was Friedman (comparison between d3 and other days in primary infection, and d1 and other time points in secondary infection). (B) Kinetic evolution of KLRG1+ and KLRG1- γδ TEM. (A-B) These experiments were repeated twice with comparable results.

Then, we assessed the presence of KLRG1 on γδ TEM, based on the reported expression of this receptor on inflationary CMV-specific CD8+ αβ T cells [32, 33]. The percentage of KLRG1+ among γδ TEM importantly increased from d0 to d92 in MCMV-target organs and blood (S1B Fig). As depicted in Fig 2B (upper panels), the γδ T cell secondary response was dominated by KLRG1+ γδ TEM in the lung, while both KLRG1- and KLRG1+ γδ TEM subsets increased in the liver. Interestingly, the kinetic evolution of KLRG1+ γδ T cell numbers in blood (Fig 2B, lower panels) resembled that of lung (Fig 2B, upper left), suggesting possible involvement of circulating γδ TEM KLRG1+ cells in the pulmonary γδ T cell response to MCMV.

### 3. MCMV leaves a long-lasting imprint on γδ T cells

To understand how previous contact with MCMV shapes γδ T lymphocytes in the long-term towards increased antiviral activity, we first conducted a comparative phenotypic analysis of γδ T cells from three months infected *versus* uninfected TCRα^-/-^ mice. As shown in S2A Fig, KLRG1 and CX3CR1 were co-expressed on γδ TEM, while KLRG1 and PD1 showed mutually exclusive expression (S2B Fig). The CX3CR1 receptor, which binds to the endothelial homing chemokine fractalkine, was previously found on the cell surface of Vδ1+ T lymphocytes induced by HCMV infection [34, 35]. In samples from MCMV-naïve mice, the majority of γδ TEM were KLRG1-CX3CR1- (Fig 3A, upper panels), akin to γδ TCM (lower panels).

**Fig 3.**
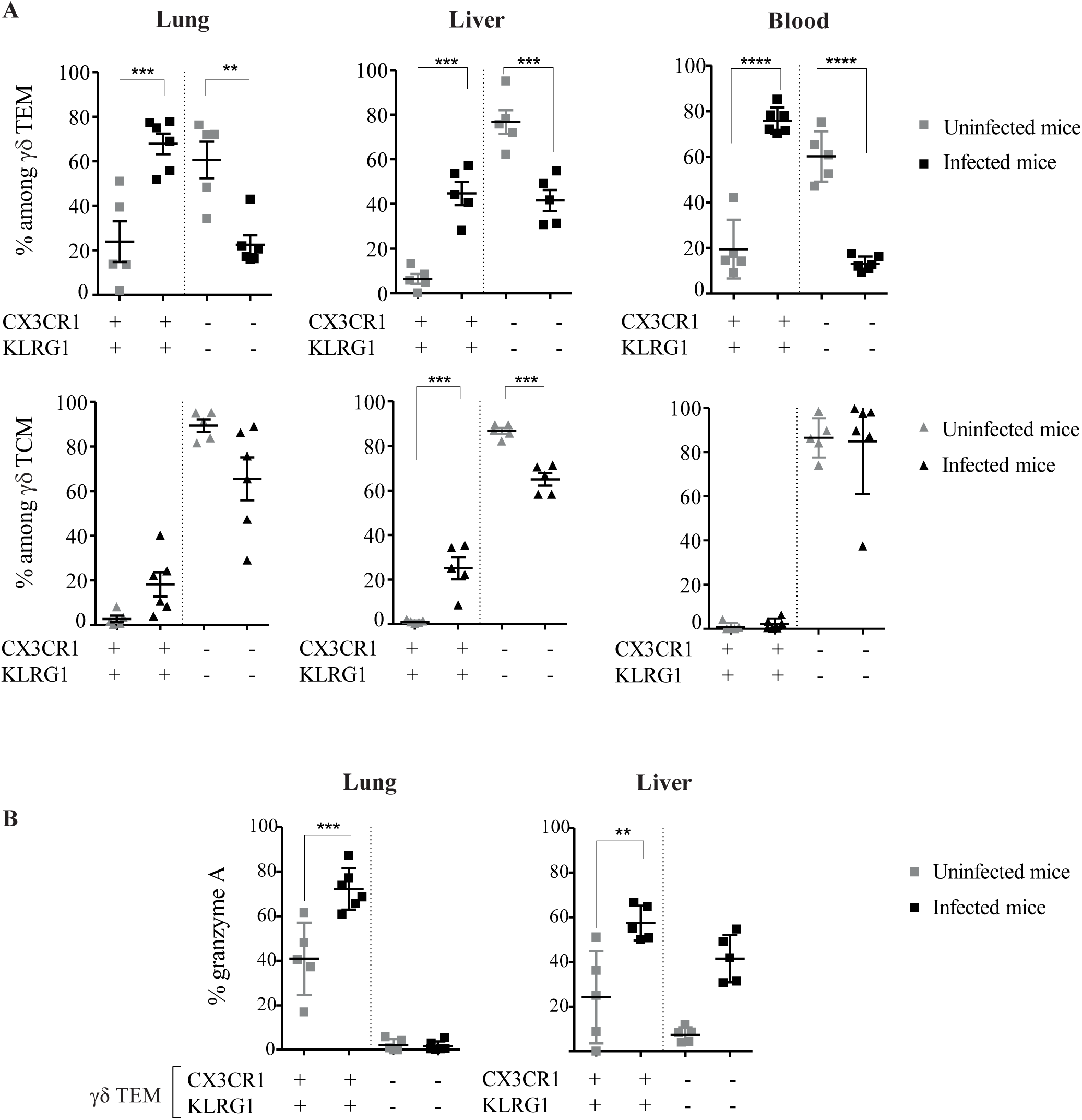
MCMV leaves a long-lasting imprint on γδ T cell phenotype. Long-term (d92) TCRα-/- MCMV infected mice (dark) were compared to age-matched uninfected TCRα-/- mice (grey). (A) Proportions of KLRG1+ CX3CR1+ and KLRG1-CX3CR1-cells among γδ TEM (upper panels) and γδ TCM (lower panels) from indicate samples in individual mice. (B) Intracellular Granzyme A expression in gated CX3CR1+KLRG1+ and CX3CR1-KLRG1-γδ TEM populations from lung, liver and blood. Data are representative of 2 experiments. Statistical test was 1-way ANOVA.

However, whereas γδ TCM remained mostly KLRG1-CX3CR1-in d92 infected mice, the proportion of KLRG1+CX3CR1+ cells among γδ TEM markedly rose upon infection, especially in blood (Fig 3A). Intracellular analysis of granzyme A further showed that KLRG1+CX3CR1+ γδ TEM from target organs of long-term infected mice exhibit higher cytolytic potential, comparatively to age-matched control mice (Fig 3B).

We next performed a multiplex gene expression analysis of immunology factors and receptors expressed by γδ T cells purified from organs of uninfected and MCMV-infected mice (S3 Fig). Among the transcripts that were increased concomitantly in the liver and lung at d92 (S3A Fig), there were genes associated with cytotoxicity (*GzmA, GzmB, Prf1*), inhibitory receptors (*Klrc*), as well as *Cx3cr1*. Evolution of these factors further showed their tendency to increase after reinfection (d7), especially in the lung (S3B Fig). In contrast, we evidenced a decrease of *Ccr7* and *Cdkn1a* (cyclin dependent kinase inhibitor 1) in long-term induced γδ T cells from liver and lung (Fig S3A Fig).

Taken together, these results demonstrate that, as observed for αβ T cells, MCMV leaves a long-lasting imprint on γδ T cell phenotype and function, characterized by a raise of cytotoxic KLRG1+CX3CR1+ cells among γδ TEM in target organs and blood.

### 4. Long-term γδ TEM expressing KLRG1/CX3CR1 preferentially localize in organ vasculature

To gain a better knowledge on the localization of these long-term effector γδ T cells likely involved in the local/tissue antiviral control, we discriminated vascular *versus* parenchyma γδ T cell memory subsets, using anti-CD45 intravascular injection. The majority of γδ T cells from the lung and liver of d92 infected mice appeared to be CD45+, thus positioned close to the vasculature (IV+) (Fig 4A). This repartition was not a consequence of infection since similar results were obtained in control mice. Interestingly, γδ TCM from infected lung were mostly found in the IV+ fraction, suggesting that these cells could be recruited to tissues from lymph nodes through blood flow (Fig 4B). CX3CR1+KLRG1+ γδ TEM were also largely enriched in the IV+ fraction (Fig 4C and 4D), in accord with preferential expression of CX3CR1 on peripheral memory T cells (TPM) [36]. Pulmonary “resident” γδ TEM cells were indeed mainly composed of CX3CR1-KLRG1-cells expressing PD1 (Fig 4D). When comparing infected and uninfected mice, our results reveal a tendency toward higher CX3CR1+KLRG1+ and lower CX3CR1-KLRG1-PD1+ γδ TEM in both IV+ and IV-cells from liver and lung.

**Fig 4.**
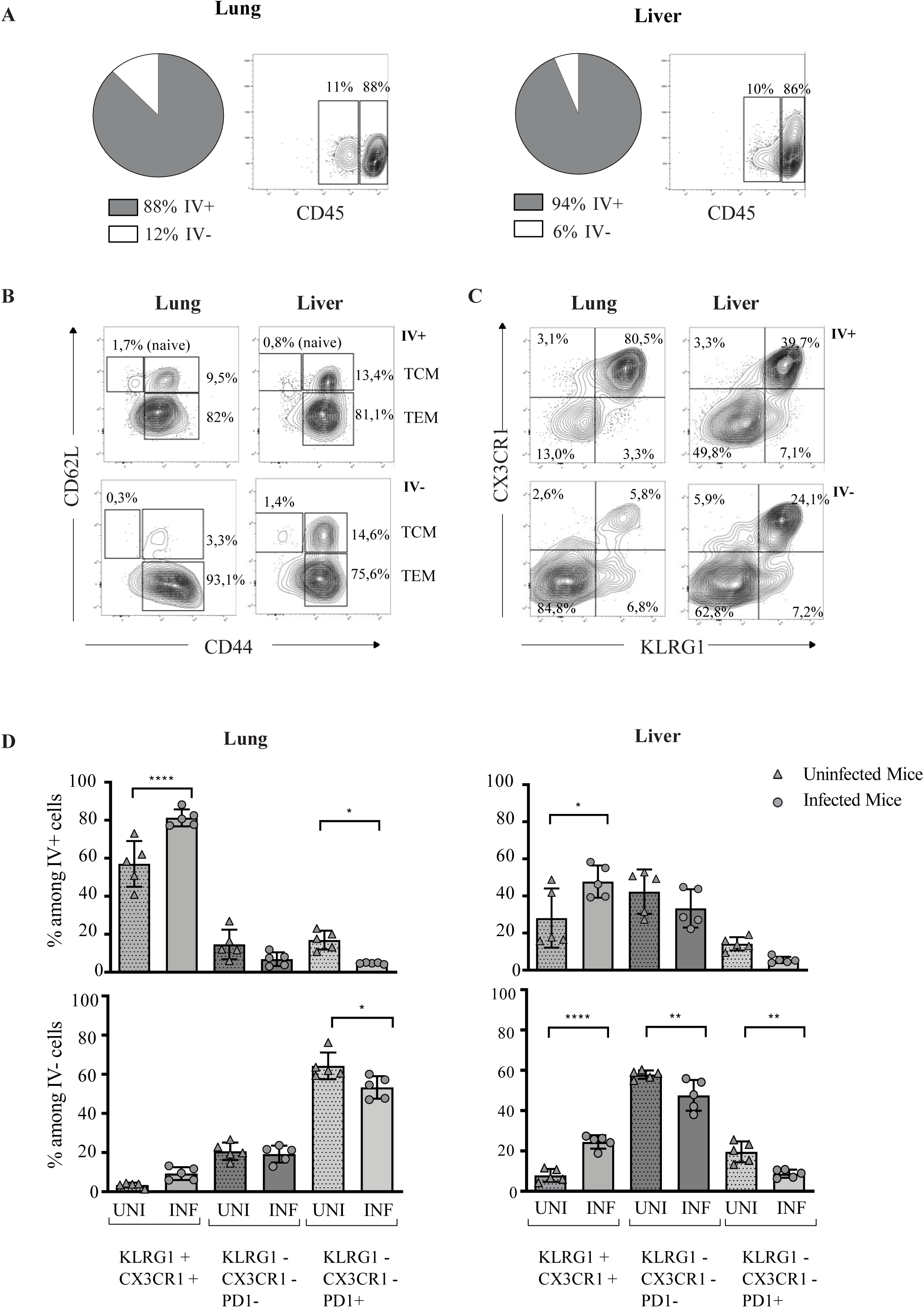
γδ T cells expressing KLRG1/CX3CR1 preferentially localize in the vasculature. Long-term MCMV infected TCRα-/- mice (INF) (n=5) or naïve age-matched control mice (UNI) (n=5) received anti-CD45 labeled mAb intravenously. Mice were sacrificed 5 min after antibody injection. (A) Repartition of γδ T cells in the intravascular positive (IV+) or intravascular negative (IV-) fraction in indicated organs from infected mice. Pie charts show the mean percentages of γδ T cells from 5 mice in each fraction. Representative contour plots obtained for one mouse out of 5 are shown. (B) Repartition of γδ naïve, TCM and TEM in the IV+ or the IV-fraction from organs of one representative infected mouse. (C) Repartition of γδ TEM subtypes within the IV+ or IV-compartment according to KLRG1 and CX3CR1 expression. (D) Comparative analyses between control uninfected (triangle) or infected (circle) mice, of the repartition of the γδ T cell memory subtypes within the IV+ or IV-compartment, according to the expression of KLRG1, CX3CR1 and PD1. Data represent the mean +/- SD of proportions of γδ T cell memory subsets from 5 infected mice or 5 naïve mice in one representative experiment out of 2. Statistical test was 1-way ANOVA.

Thus, long-term local/tissue γδ T cell response to MCMV infection probably involves cells in both vascular and parenchyma compartments, but the most prevalent subset (i.e. γδ TEM co-expressing KLRG1 and CX3CR1) preferentially localize in the vasculature.

### 5. Temporally Blocking γδ TCM egress from lymph nodes poorly affects viral load control in organs from long-term infected mice

Our next goal was to determine whether blocking lymph nodes egress of TCM would affect the long-term γδ T cell response to MCMV. To do so, we used fingolimod (FTY720) and analysed the consequence of short-term (11 days) treatment on γδ TCM/TEM numbers and viral control. An important decrease of γδ TCM counts was evidenced in blood, lung and liver when comparing untreated and FTY720-treated TCRα^-/-^ mice (Fig 5A, upper panels). In contrast, γδ TEM numbers remained constant in organs from treated mice, although they were reduced in blood (Fig 5A, lower panel). FTY720-treatment poorly affected blood poorly affected the control of viral loads in long-term infected TCRα^-/-^ mice, since no statistical differences were evidenced in organs from control *versus* FTY720-treated mice (Fig 5B). These results suggest that long-term γδ T cell response to MCMV involves tissue control of viral loads by local γδ TEM.

**Fig 5.**
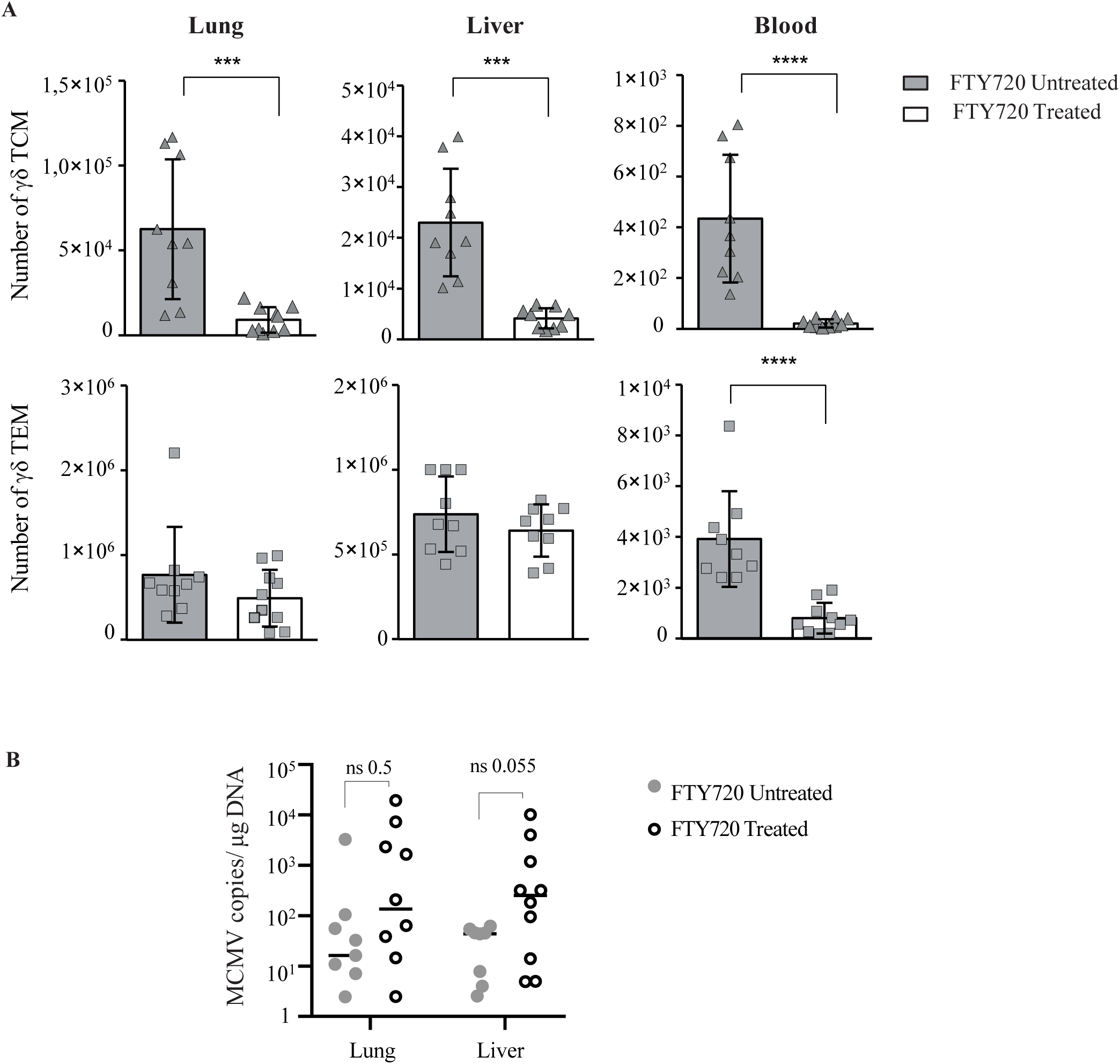
Blocking γδ TCM egress from peripheral organs poorly affects viral load control in long-term infected mice. Long-term MCMV infected TCRα-/- mice (n=10) were treated (n=5) or not (n=5) with FTY720 for 11 days before sacrifice. (A) Histograms show the number of γδ TCM (upper panels) and TEM (lower panels) in organs (total numbers) or blood (per 25 µl). (B) Viral loads in organs of treated or untreated mice were quantified by qPCR. Each point indicates one individual mouse and represents MCMV copy numbers in 1 µg of total DNA. Data are pooled from two independent experiments (n = 5 per group). (A-B) Error bars represent the standard error of the mean of γδ T cell numbers (A) and of the median of MCMV copy numbers (B). Differences were evaluated by the Mann-Whitney statistical test.

### 6. Viral control after reinfection engages γδ-TCR signaling

To test whether the memory antiviral potential of γδ T lymphocytes was dependent on TCR signaling, we used the anti-γδ TCR mAb clone GL3, which induces down regulation of the surface expression of the TCR, rendering γδ T cells invisible and poor responders to TCR-specific stimulation [37]. Three months post primary MCMV infection, TCRα^-/-^ mice received intravenous injection of GL3 or isotypic control mAb, followed (24h later) by MCMV secondary challenge (Fig 6A). The effect of the anti-γδ TCR mAb treatment was confirmed by an absence of TCRδ surface expression (Fig 6B). Seven days post-re-infection, a substantial increase of viral loads was observed in organs from mice that had received the anti-γδ TCR mAb comparatively to control mice (Fig 6C) indicating TCR involvement in γδ T cell control of CMV.

**Fig 6.**
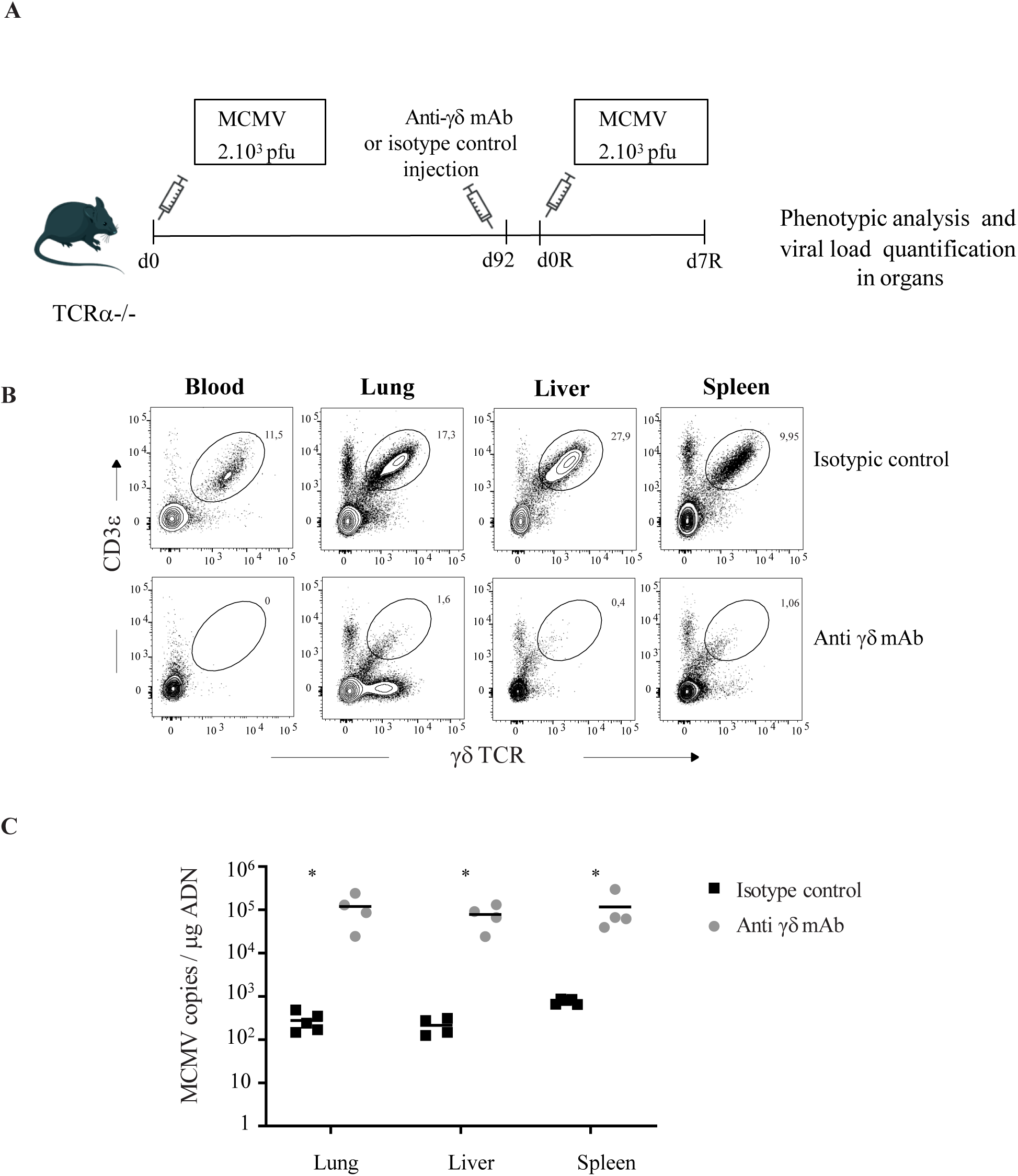
Viral control after reinfection engages γδ-TCR signaling. TCRα-/- mice (n=10) were infected with MCMV (2.10^3^ PFU) at day 0. At day 92, mice received isotype control or anti-γδ mAb (n=5). 24h after, mice were re-challenged with similar doses of MCMV. (A) Experimental scheme. (B) Staining of CD3ɛ+/ γδ+ cells in mice treated with anti-γδ TCR or control mAb at day 7 post reinfection. (C) Viral loads/1 µg of total DNA for individual mouse were quantified in organs by qPCR 7 days post-reinfection. Statistical tests were Mann Whitney. The experiment was repeated twice with comparable results.

### 7. MCMV drives the expansion of a private and adaptive-like TCRδ repertoire

As the γδ TCR is involved in the viral control by long-term-induced γδ T cells in our model, we tested whether there were changes in the TRD (TCRδ chain) repertoire upon MCMV infection. We performed TRD CDR3 NGS analysis on blood at different time points post-primary (short-term (d21), long-term (d80)) and post-secondary (d7R) MCMV infection. At d7R, the TRD repertoires of lung, liver and spleen were analyzed in parallel. Age-matched control mice were studied simultaneously to take into account possible impact of ageing [38]. No major differences were observed between uninfected (n=3) and infected mice (n=5) in TRDV usage, CDR3 length distribution and number of N additions (S4A Fig).

While the TRD repertoire appeared to be relatively stable between d21 and d80, important changes were observed after infection (d21) and reinfection (d7R) (Fig 7A, upper panels). Some clonotypes present at d0 (dark blue, mice 1 and 2) were barely found at d21, leaving space to other clonotypes (see for example mouse 1, where clone CALMEREAFWGGELSATDKLVF (TRDV2-2) became highly prominent). The secondary γδ T cell response to MCMV was composed by a combination of clonotypes expanded post-primary infection and, more unexpectedly, of novel expanded clonotypes (for example mouse 2 and 3, Fig 7A in orange). Note that at d7R a higher TRD blood repertoire overlap with organs was observed compared to d0 (S5 Fig), indicating that the blood γδ T cell response to MCMV reflects (at least partially) what occurs in organs at a given time point.

**Fig 7.**
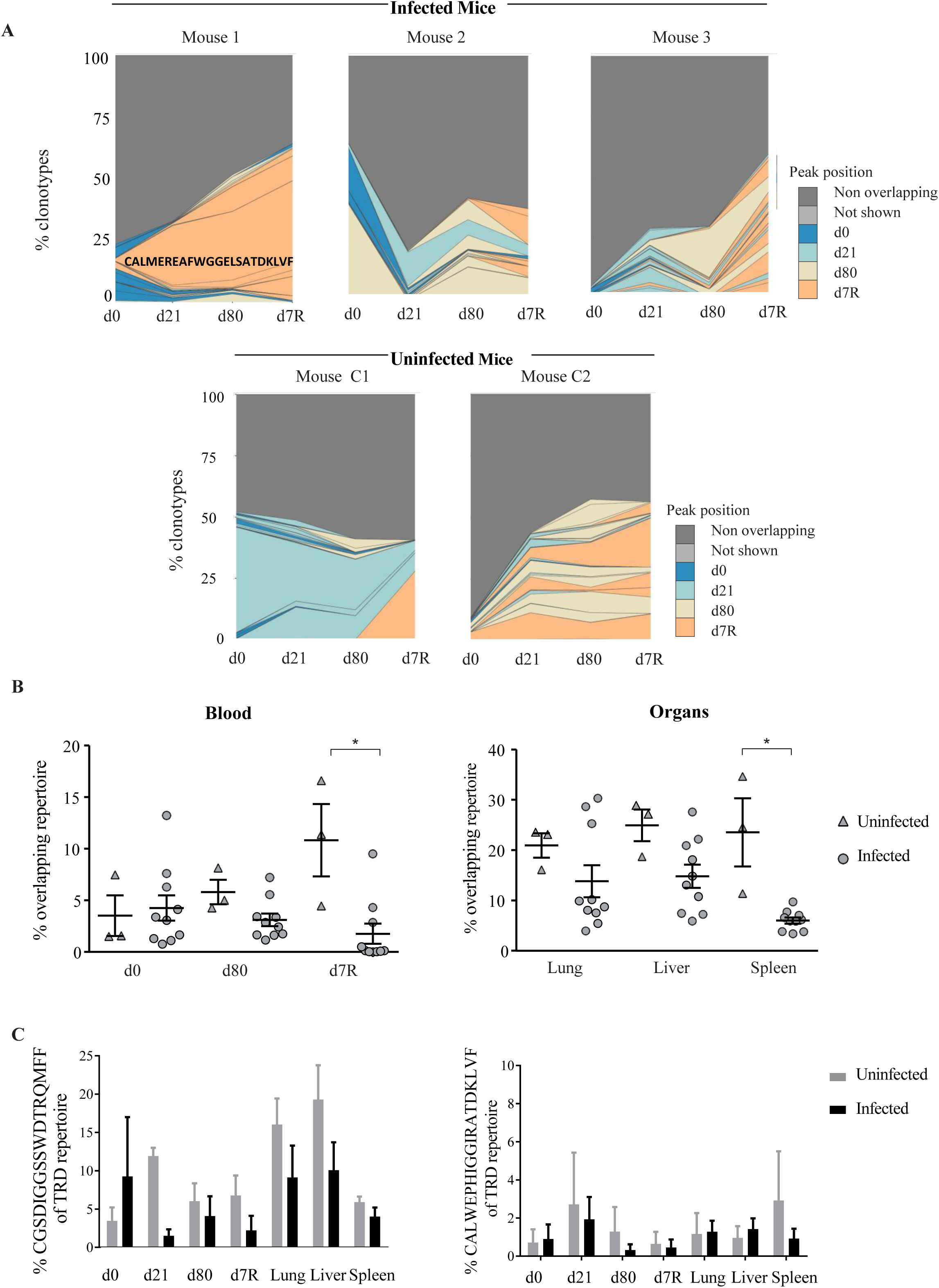
Expansion of private CDR3δ repertoire following MCMV infection. TCRα-/- mice (n=5) were infected with MCMV. Age matched uninfected TCRα-/- mice (n=3) were used as controls. Blood was drawn at different time intervals post-infection and at d7 post-reinfection. (A) Examples of clonotype tracking stackplots for 3 mice in the infected group and 2 mice in the control group. Detailed profiles for top 100 clonotypes, as well as collapsed (“Not-shown” in dark gray) and non-overlapping (light gray) clonotypes found in blood of MCMV-infected mice (Upper panels) or age-matched control mice (Lower panels). Clonotypes are colored by the peak position of their abundance profile. (B) Geometric mean of relative overlap frequencies (F metrics) within pairs of blood (left) or of organ (right) samples, each dot represents the F value (x100) of a pair of samples. Statistical tests were Mann Whitney. (C) Percentage of CGSDIGGSSWDTRQMFF (left), and CALWEPHIGGIRATDKLVF (right) sequences in TRD repertoire, in blood and organs from infected and naïve age matched control mice. Statistical test was 1-way ANOVA.

In comparison to infected mice, the kinetic evolution of clonotypes in blood of uninfected mice remained more stable overall, although some changes also occurred consecutively to control medium injection (Fig 7A, lower panels). Notably however, the frequency of shared clonotypes between individuals of the infected mice group (i.e. F value, a repertoire overlap measure) was lower than that of the control group (Fig 7B), while no clear influence could be observed on the diversity of the repertoire (S4B Fig), indicating that the TRD repertoire became more private upon MCMV infection (unique to each individual). Among the public sequences highly shared between samples, we identified the innate CDR3 amino acid sequences CGSDIGGSSWDTRQMFF (*TRDV4* sequence) [39–42] and CALWEPHIGGIRATDKLVF (TRAV15-1-DV6-1 sequence) related to so-called NK T γδ [43, 44] (for nomenclature see [45]). In line with an increased private repertoire upon infection, a tendency towards lower representation of these shared clonotypes in infected *versus* uninfected mice was observed (Fig 7C).

In sum, our results highlight the preferential amplification and/or recruitment of a private and adaptive-like TRD repertoire upon MCMV infection. The γδ T cell secondary response to MCMV appears to involve both γδ TCR clonotypes expanded after the primary MCMV encounter as well as novel expanded clonotypes.

### 8. Long-term MCMV-induced γδ T cells confer protection to T cell deficient hosts upon adoptive transfer

To confirm the protective antiviral function of long-term induced γδ T cells, γδ T cells were isolated from the spleen of three month-MCMV-infected TCRα^-/-^ infected mice, and transferred into CD3ε^-/-^ hosts (Fig 8A). γδ T cells isolated from MCMV-infected mice importantly increased the survival rate of CD3ε^-/-^ hosts that are infected with MCMV one day after the transfer (Fig 8B, left). Improved viral control in γδ-T cell bearing versus T-cell deficient mice was evidenced by lower viral loads in organs (Fig 8B, right). In a second set of experiments, γδ-T cells were sorted from age-matched control mice (5 months old) and transferred into T-cell deficient hosts. A control group of mice received γδ T cells from d92 MCMV-infected mice as in the first experiment. CD3ε^-/-^ hosts bearing “naïve” γδ T cells (i.e. unprimed by MCMV) became moribund within same time-delay as T cell deficient hosts and bared comparable viral loads at sacrifice, in contrast to CD3ε^-/-^ bearing MCMV-primed γδ T cells (Fig 8C, upper panel). Higher quantities of transaminases were found in age-matched naïve γδ -T cell bearing mice, comparatively to CD3ε^-/-^ hosts that had received d92 MCMV-primed γδ T cells (Fig 8C, lower left panel). Alongside, mice transferred with unprimed γδ T lymphocytes had lower γδ T cell counts (Fig 8C, lower right panel). Finally, we analyzed γδ T cells in 5 CD3ε^-/-^ mice that had survived for 130 days (i.e. transferred with d92 MCMV-primed γδ T cells). The repartition of γδ T cell memory subtypes in γδ-bearing CD3ε^-/-^ and d92 infected TCRα^-/-^ mice was comparable, evidencing efficient reconstitution (S6 Fig).

**Fig 8.**
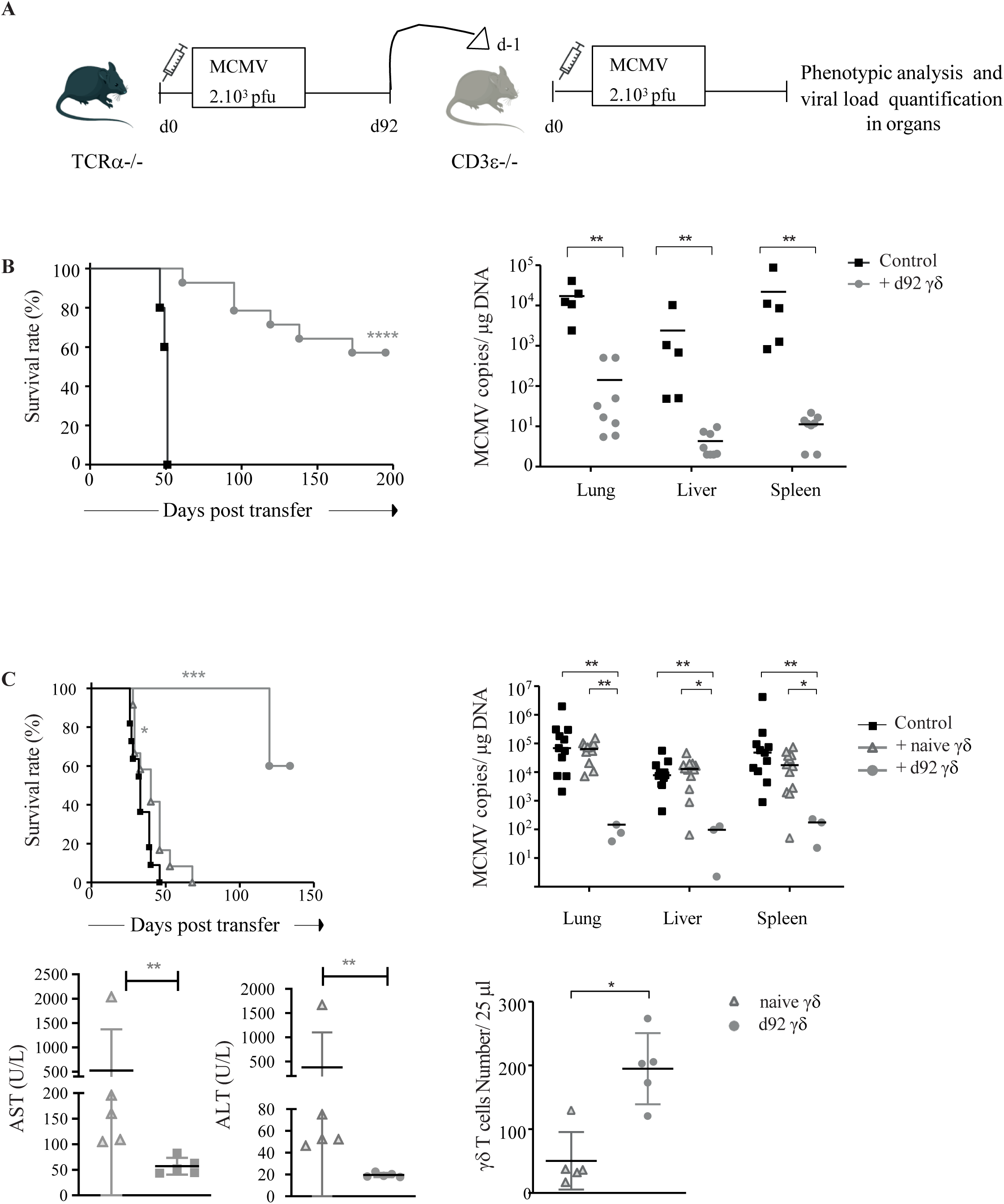
Long-term induced γδ T cells confer protection to T cell deficient hosts upon adoptive transfer. (A) Experimental scheme. γδ T cells were sorted from the spleen (purity > 95%) of 3 months infected TCRα-/- mice and 1.10^6^ cells were transferred into CD3ɛ-/- mice. The day after, CD3ɛ-/- hosts were infected with MCMV (2.10^3^ PFU). (B, left) Survival curve of mice given long-term MCMV-primed γδ T cells (grey line, n=14) and of CD3ɛ-/- control mice (dark line, n=5). Mice were sacrificed when losing 10% weight. (B, right) Quantification of viral loads by qPCR, in organs from 5 T-cell deficient mice at sacrifice (dark squares) and 8 γδ-bearing mice that had survived until d130 (grey circles). (C) 1.10^6^ γδ T cells from age-matched TCRα-/- control mice were transferred into CD3ɛ-/- mice that were MCMV-infected the day after. (C, upper left) Survival curve of mice given naïve γδ (n=12), MCMV-primed γδ (n=5), and of untransferred CD3ɛ-/- mice (n=11). (C, upper right) Quantification of viral loads by qPCR, in organs from CD3ɛ-/- control mice (n=11), naïve γδ T cell bearing mice (n=11), and MCMV primed γδ T cell bearing mice (n=3) (C, lower panels) Serum quantification of aspartate aminotransferase (ALT) and alanine aminotransferase AST (left) and number of γδ T cells in blood (right) at day 30 post transfer, in 5 mice receiving naïve or MCMV-primed γδ T cells. Statistical tests were Mann Whitney. Log-rank test were used for Kaplan-Meier survival curves. Survival curves and quantification of viral loads were from one representative experiment out of two.

Thus, previous contact with MCMV endows γδ T cells with an enhanced protective potential which may (at least partially) depend on a longer half-life in T-cell deficient hosts.

### 9. The protective function of d92 MCMV-induced γδ T cells from the spleen principally relies on the presence of γδ TCM

Long-lasting memory has long been attributed to αβ TCM, while the function of γδ TCM remains poorly studied. At day 92 post-MCMV infection, γδ T cells from the spleen of TCRα^-/-^ mice comprised CD44+CD62L+ (TCM), CD44+CD62L- (TEM) KLRG1+ (mostly CX3CR1+) and TEM KLRG1- (mostly CX3CR1-) (S6 Fig on the top right). To gain knowledge on the protective function of these memory γδ T cell subsets, we sorted γδ TCM, TEM KLRG1+ and TEM KLRG1-from the spleen of d92 MCMV-infected TCRα^-/-^ mice, and their protective function after transfer was assessed as above. Interestingly, γδ TCM was the only subtype able to confer good protection to T-cell deficient hosts upon adoptive transfer (Fig 9A, upper panel). In moribund mice from the TEM KLRG1+ and KLRG1-transfer groups, viral loads (Fig 9A, lower panels) and transaminase levels (Fig 9B) were elevated, with no statistical difference comparatively to the non-transferred group. In contrast, for both readouts, significant differences were observed between CD3ε^-/-^ control mice and surviving mice from the TCM transfer group (Fig 9A, lower panels and fig 9B). Markedly, the majority of γδ T cells recovered in γδ TCM recipient mice display a CD44+CD62L-TEM phenotype, and were for the most composed of KLRG1+CX3CR1+ cells in lung, and KLRG1-CX3CR1-cells in the liver and spleen (Fig 9C). Finally, we showed that γδ TCM from control and long-term MCMV-infected mice exhibit higher proliferative potential in culture with IL15, comparatively to γδ TEM (S7 Fig).

**Fig 9.**
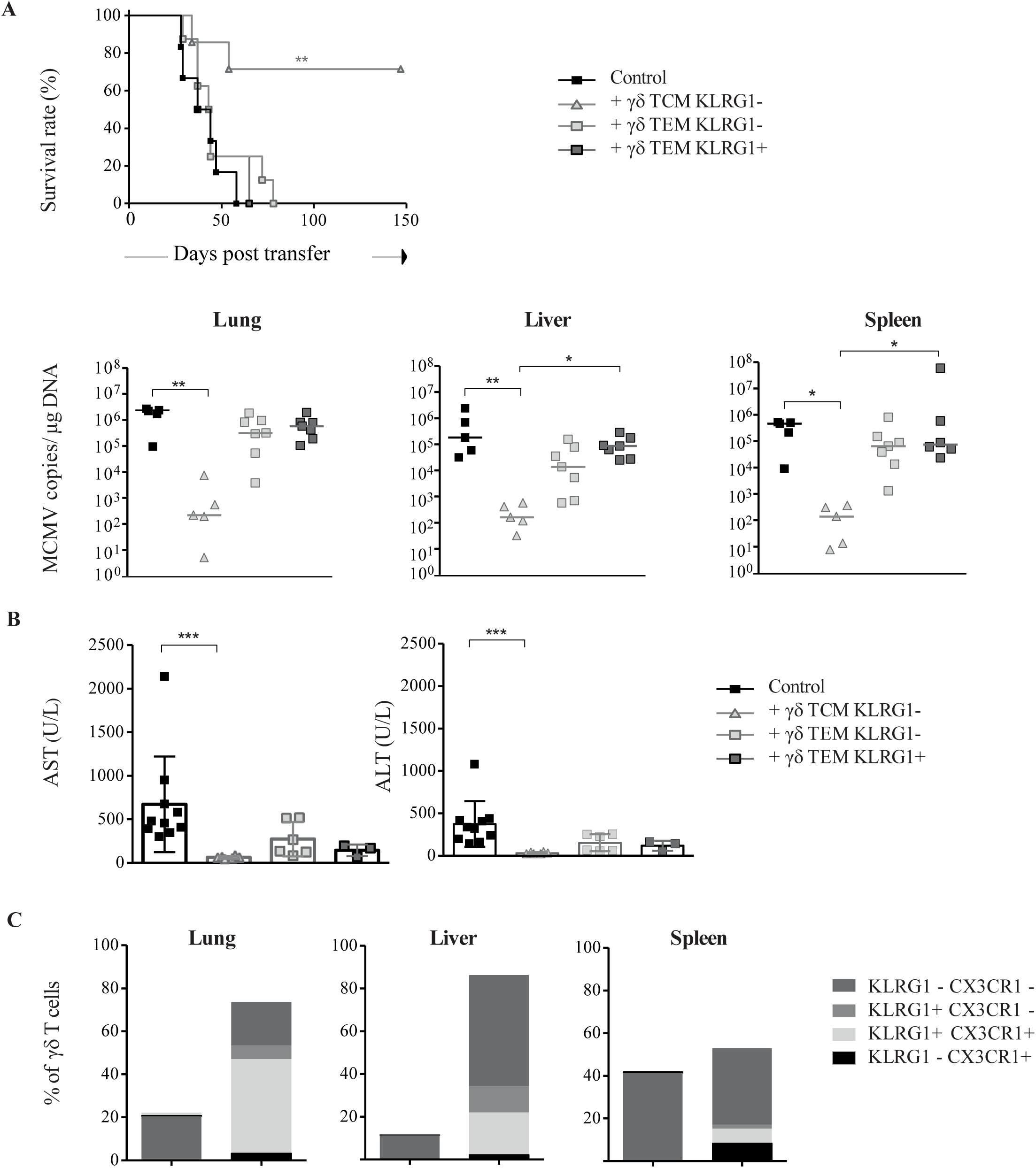
The protective function of d92 MCMV-induced γδ T cells from the spleen principally relies on the presence of γδ TCM. (A, upper panel)) Survival curve of CD3ɛ-/- mice given 200000 γδ TCM KLRG1- (n=7), γδ TEM KLRG1- (n=8) or γδ TEM KLRG1+ (n=8). The experiment was repeated twice with comparative results. (A, lower panel) Quantification of viral loads by qPCR in organs from recipient mice at sacrifice. (B) Serum quantification of ALT and AST. Mice given γδ TEM KLRG1- and KLRG1+ cells were sacrificed when losing 10% weight, and mice given γδ TCM KLRG1-were sacrificed at the end of the experiment, at d150 (n=5). (C) Distribution of γδ T cell memory subsets in organs from γδ TCM recipient mice that had survived (mean numbers). Same statistical tests are as in Fig 8.

These results univocally show that γδ TCM are precursors of KLRG1+ and KLRG1-γδ TEM. Low numbers of MCMV-primed γδ TCM are sufficient to maintain long-term antiviral activity against MCMV in T-cell deficient hosts, possibly through their capacity to survive and to generate γδ TEM able to control viral loads in target organs.

## DISCUSSION

In contrast to increasing numbers of studies showing the memory potential of γδ T cells during acute resolving infections, data describing γδ T cell memory in persistent infections are missing. CMV is the prototypical β-herpesvirus establishing lifelong latent infection. Anti-CMV memory responses are essential to control viral reactivation and/or reinfection events that commonly take place in SOT. The role of γδ T cells could be significant in this context where immunosuppressive/ablative treatments cause suboptimal and/or delayed αβ T cell responses [22]. In the present study, we used subsequent MCMV infections of TCRα^-/-^ mice to decipher the memory potential of γδ T cells against CMV. Our data depict a more rapid and more efficient γδ T cell secondary *versus* primary antiviral response in MCMV-target organs and blood, with the implication of γδ TEM. They are in line with the faster increase of non-Vγ9Vδ2 TEMRA cells in blood from secondary infected HCMV-seropositive *versus* primary infected HCMV-seronegative renal transplant recipients, which associates to shorter infection resolution [26, 29]. Stressing the value of our mouse model, our study points out that γδ T cell memory function can operate independently of priming with αβ CD4 T cells. Furthermore, our adoptive transfer experiments show a major role for γδ TCM in the maintenance of long-term antiviral activity.

Despite long-term survival of TCRα^-/-^ infected mice, DNA copies of MCMV were still detected in the liver and lung at d92. This suggests the continuous production of infectious virus in the absence of αβ T cells. Even so, MCMV drove the accumulation of γδ TEM that were mostly PD-1^neg^, unlike T cells responding to chronic viruses such as non-cytopathic LCMV clone 13 [7]. In fact, γδ TEM from d92-infected mice expressed KLRG1, a typical feature of CMV-induced αβ CD8 TEM that was associated to antigen persistence rather than to viral replication [46, 47]. These results emphasize the importance of the intrinsic nature of pathogen in determining T cell fate, although the route and dose of infection may also operate [48, 49].

We show here mutually exclusive expression of KLRG1 and PD-1 on γδ TEM, related to their anatomical localization. KLRG1+ γδ TEM co-expressed CX3CR1 and were mainly located in the blood and intravascular compartment from organs, while PD-1+ γδ TEM prevailed beyond the vasculature. Our results are in line with the recent study by Oxenius’s team, who depicted a rise of pulmonary intravascular (IV+) KLRG1+ αβ CD8 TEM driven by MCMV [50]. Interestingly, the increase of KLRG1+ γδ TEM in the intravascular compartment occurred concomitantly to a decrease of KLRG1-γδ T cells in the IV-fraction, suggesting a possible modulation of this receptor while γδ T cells transit from the parenchyma to the blood circulation, and reciprocally. The accumulation of KLRG1+ γδ TEM in long-term infected mice might thus reflect their continuous dissemination through the bloodstream in response to systemic MCMV infection.

Priming of γδ T cells with MCMV induced transcription of genes associated to cytotoxicity, in accord with their reported capacity to kill CMV-infected fibroblasts, a function that could be important to control the virus *in vivo* [28]. In the liver, where the virus early disseminates *via* the intraperitoneal route, the fast mobilization of KLRG1-Granzyme A+ cells after secondary viral challenge sustains their implication in the control of local infection, possibly through killing of MCMV-infected parenchymal cells (i.e. hepatocytes). Long-term MCMV-primed γδ T cells also upregulated genes encoding NKR, including signaling lymphocyte activation molecules (SLAM). The latter are commonly found on cytotoxic effectors and may act as important rheostats to fine-tune their functions. The modulation of these transcripts along the course of MCMV infection and reinfection is reminiscent to repetitive antigen exposure [51], suggesting continuous triggering of the γδ TCR by still-unknown MCMV-induced antigens. The concomitant downregulation of genes/protein involved in trafficking into lymph nodes suggest a preferential function of these cells in the tissues such as the liver and lung.

In spite of applying a second boost of MCMV, viral loads in organs remained mostly stable or even decreased after reinfection, although interindividual variations were noticed. Our results suggest that rapid virus handling upon MCMV secondary challenge might benefit from the long-lasting molecular imprint left on γδ T cells by first MCMV encounter. This hypothesis is supported by the capacity of long-term MCMV-primed γδ T cells to confer protection upon adoptive transfer into T-cell deficient hosts; in contrast, the time-delay needed to proliferate and differentiate probably precludes MCMV-naïve γδ T cells to counteract the amplification and dissemination of viral copies. d92-induced γδ T cells might have a higher survival rate, proliferative capacity, and/or effector function than unprimed γδ T cells. Improved survivability of viral-primed γδ T cells is suggested by the decline of transcripts encoding the cell-cycle inhibitor p21 (*Cdkn1a*), and by their maintenance upon transfer into T cell deficient hosts.

First evidence for γδ T cell memory responses were put forward in the early 2000 and implicated phosphoantigen-reactive Vγ9Vδ2 in primates [52–54]. In mice, pioneering work was carried out by Lefrançois’s team who showed higher increase of Vγ6Vδ1 cells in mesenteric lymph nodes after *Listeria monocytogenes* (*Lm*) secondary and even tertiary oral challenge, relatively to primary infection [55, 56]. Since then, increasing numbers of studies have shown the memory potential of γδ T cells in infections ([41, 42, 44] and reviewed in [13–15]). The γδ T cell subtypes described in most of these studies are pre-activated lymphocytes with innate-like features and limited TCR diversity. In contrast, a tendency towards an increase in TRD diversity could be noticed in blood after MCMV reinfection, concomitant to a decrease of the highly frequent innate like CGSDIGGSSWDTRQMFF sequence, and to the acquisition of a more private TRD repertoire. Since innate-like γδ T cells are commonly shared between mice, our results suggest the mobilization/amplification of private adaptive-like γδ T cell clones soon after MCMV encounter. They are in line with our previous data showing, by the use of CD3ɛ^-/-^ mice receiving TCRα^-/-^ bone marrow graft, that fetal-derived γδ T cells are dispensable for long-term MCMV protection in the adult mice [27]. The participation of innate-like γδ T cells in the antiviral response remains possible, and might be particularly relevant in humans where congenital HCMV infection can occur [34, 57]. Interestingly, expansion of private CDR3β clones at the expense of more public ones was described by Friedman and colleagues following immunization with either self or non-self MHC-restricted peptides [58].

The implication of adaptive-like γδ T lymphocytes in the antiviral response is supported by the rise of the γδ TEM/TCM ratio along the course of MCMV infection. Likewise, HCMV infection induces the differentiation of non-Vγ9Vδ2 T cells from a naïve to TEMRA phenotype [29, 59]. However, in contrast to the studies in humans which describe HCMV as a major driver of TRD repertoire focusing over time in transplant patients [22, 35, 60] the overall TRD diversity in blood from long-term infected mice (d80) was similar to that of control mice. This suggests that the TRD repertoire somehow stabilizes at distance from MCMV challenge, likely due to the loss of some adaptive-like γδ T cell effectors, and to the high frequency of innate like γδ TEM in mice regardless of infection. Alternatively, the transplant setting in the HCMV studies may contribute to the high expansion of particular γδ clonotypes. Importantly, TCRγδ signaling was essential to afford viral load control after reinfection. Whether γδ T cell memory response involves TCR binding of diverse MCMV-induced antigens remains to be elucidated [16]; [61]. Counterintuitively, γδ clonotypes that dominated the chronic phase of infection (d80) were not necessarily expanded in the initial phase of infection (d21). Likewise, expansion of γδ T cell clonotypes after first MCMV challenge was not a prerequisite for their involvement in the secondary response. In an elegant study, the group of Buchholz recently showed that, during MCMV infection, long-term immunodominance of a single-cell derived αβ T cell family is not determined by its initial expansion, but is rather predicted by its early content of T central memory precursors [62]. Moreover, inflationary KLRG1+ αβ CD8 T cells have a reduced half-life and are supposed to be maintained by continuous replenishment from early primed KLRG1-αβ T cells, among which αβ TCM expressing Tcf1 [33, 63–66]. We hypothesize that γδ TCM primed by a first MCMV encounter play a crucial role in the long-term anti-CMV response, through their capacity to proliferate and to give rise to antiviral effectors that eventually localize in target tissues or recirculate. This would explain why blocking γδ TCM egress from lymph nodes does not affect viral loads control, at least in the short term. In accordance with our hypothesis, we showed the high proliferative potential of γδ TCM *in vitro*, as well as their protective function and their ability to generate KLRG1+ γδ TEM upon adoptive transfer into T cell deficient hosts. Applying a second intraperitoneal injection of MCMV to primarily infected mice probably leads to the mobilization of novel γδ T cell responders, thus explaining the appearance of novel γδ TCR clonotypes, and to an enhanced γδ T cell secondary response.

To conclude, our results uncover memory properties of TCM γδ T cells in the context of a chronic viral infection. These data reveal the interest of investigating and targeting this subset of unconventional T cells in strategies aiming at improving antiviral vaccination approaches. TCM γδ T cells could compensate for the defect of αβ T cells in immunosuppressed individuals and are thus of particular relevance in the context of organ or hematopoietic stem cell transplantation.

## MATERIALS AND METHODS

### Mice

Specific Pathogen-free, 8-12 weeks old, C57BL/6 CD3ε^-/-^[67] as well as TCRα^-/-^ [68] were purchased from the CDTA (Centre de Distribution, Typage et Archivage Animal, Orléans, France). Experiments were performed in an appropriate biosafety level 2 facility in compliance with governmental and institutional guidelines (Animalerie spécialisée A2, Université Bordeaux Segalen, France, approval n° B33-063-916). This study was carried out in accordance with the Ethics Review Committee of Bordeaux (2016092917471799).

### Virus

MCMV (Smith strain, ATCC VR-194) was obtained from the American Type Culture Collection and propagated into BALBc mice (BALBcBy/J, Charles Rivers laboratory, Larbresle, France) as previously described. Three weeks after MCMV infection, salivary glands were collected from infected mice and used as an MCMV stock solution. Determination of virus titers was defined by standard plaque forming assay on monolayers of mouse embryonic fibroblasts (MEF). Infections were performed by intraperitoneal (i.p.) administration of desired quantity of PFU from the salivary gland viral stock.

### Flow cytometry

For lung preparation, collagenase I (50 μg/ml; Sigma) and DNase I (50 μg/ml; Sigma) were used for 1h at 37°C. Single-cell suspension from spleen, liver and enzyme treated lung were prepared by meshing organs through 70 μm nylon cell strainers in sRPMI-1640 with 8% FBS (HyClone Laboratories, GE Healthcare, Logan, Utah). After red blood cells lysis with NH4Cl (liver) or ACK (lung), lymphocytes were isolated by a discontinuous 40/80% Percoll gradient (GE Healthcare).

For Flow cutometry analysis, before antibody staining, Fc-receptors were blocked with CD16/CD32 FcR antibody (clone 93, eBioscience™). Live/dead discrimination was performed using Fixable Viability Stain 700 (BD Horizon™) according to manufacturer’s instructions.

The following monoclonal antibodies were used:

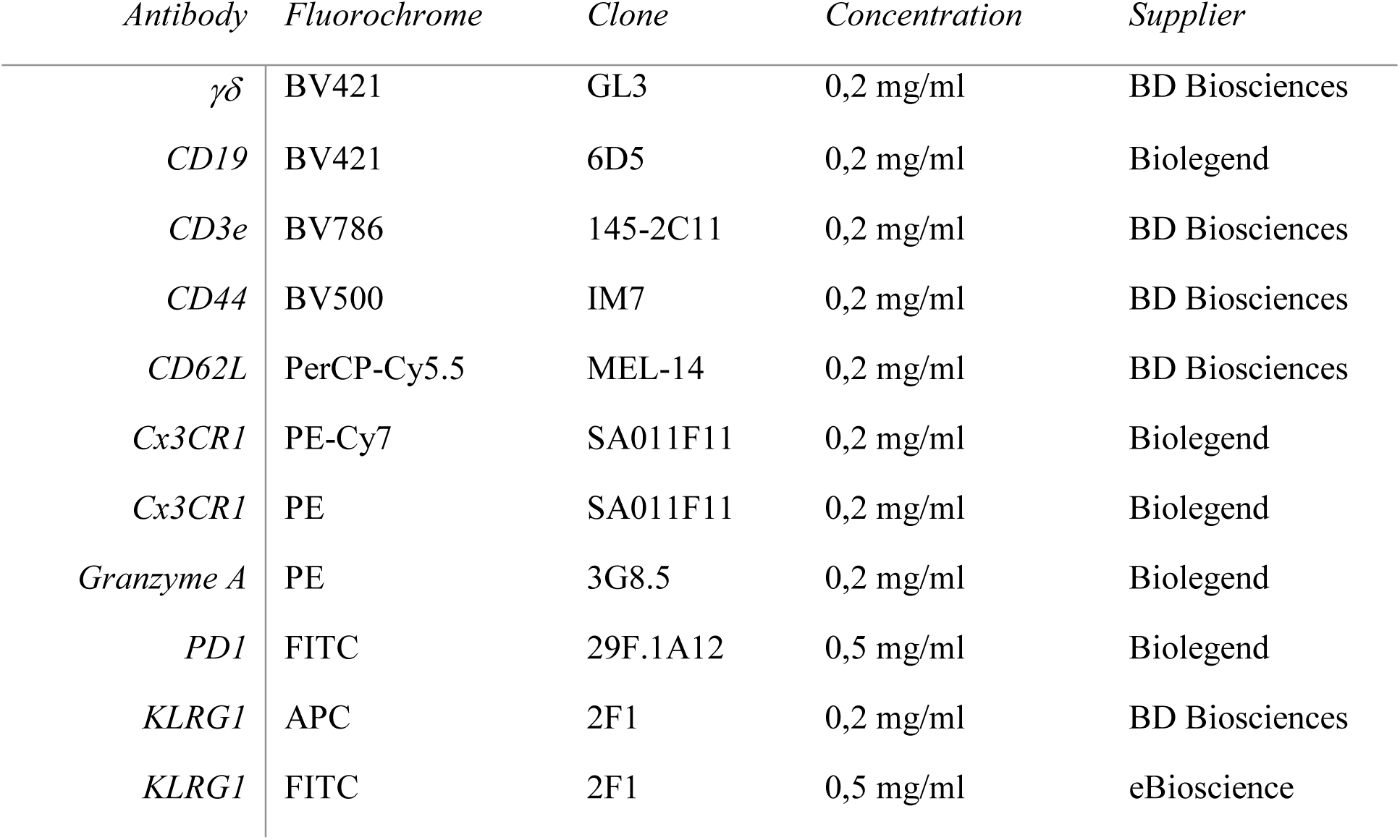

For detection of intracellular cytokines, we employed the BD Cytofix/Cytoperm™ fiixation/Permeablization kit. Cells were acquired using a LSRFortessa (BD Biosciences), and analyzed using FlowJo software (Tree Star).

### DNA extraction and quantification of MCMV DNA copy number

Genomic DNA from organs was isolated using a Nucleospin tissue purification kit (Macherey Nagel). MCMV-DNA was quantified by real-time PCR using BIO-RAD CFX with GoTaq qPCR Master Mix (Promega) and primers specific for MCMV glycoprotein B (gB) (forward primer: GGTAAGGCGTGGACTAGCGAT and reverse primer: CTAGCTGTTTTAACGCGCGG). Samples were distributed by Eppendorf epMotion 5073 automated pipetting. PCR condition was as follows: 95°C for 10 min, denaturation at 95°C for 15 s, and annealing/extension at 60°C for 1 min. Known quantities of plasmid comprising MCMV gB were used for the titration curve.

### Adoptive transfer of sorted γδ T cells

γδ T cells were sorted from the spleen of MCMV infected or control TCRα^-/-^ mice using the TCR γδ T cell Isolation kit (Miltenyi Biotec). For γδ TCM and TEM sorting, purified γδ T cells were stained and isolated using FACSAria II Sorter (BD Biosciences). Purity was > 95%. Recipient CD3ε^-/-^ mice received intravenous (i.v.) injections of a defined number of total γδ T cells or of memory γδ T cell populations. These animals were infected with 2.10^3^ pfu of MCMV the day after and followed twice a week. Mice were sacrificed when losing 10 % of their original weight or when showing signs of distress.

### In vivo treatment with anti-γδ TCR mAb

At day 92 post MCMV infection, TCRα^-/-^ mice received 200 μg/mouse of purified Hamster anti-mouse γδ T-Cell Receptor (Clone GL3) mAb, or Hamster IgG2, κ Isotype Control (Clone B81). Treatment efficiency was tested by FACS analysis in blood and organs.

### AST and ALT quantifications

Mice were bled via the retro-orbital sinus after anesthesia. Serums were collected and frozen before quantification using a clinical chemistry analyzer (Horiba Pentra C400).

### Anti-CD45 Intravascular staining

For in vivo antibody labelling, a total of 10 μg/100 μl of anti-CD45 APC (clone 30-F11 from eBioscience; 0,2 mg/ml) was injected i.v to anesthetized mice via the retro-orbital venous plexus as previously described [69]. Antibody was allowed to circulate for 5 min to label cell in circulation or closely related to the vascular space. After this lapse-time, animals were sacrificed. Spleen, liver and lung were removed. Cells were isolated for ex vivo labelling before flow cytometry.

### FTY720 Treatment

To prevent lymphocyte recirculation and egress from lymph nodes, the S1P1R agonist, FTY720 (2-amino-2-[2-(4-octylphenyl)ethyl]-1,3-propanediol) was used. FTY720 (Sigma-Aldrich) was reconstituted in ethanol and diluted in 2% β-cyclodextrin (Sigma-Aldrich) for injections. Long-term MCMV infected TCRα^-/-^ mice (d92 post primary infection) received every other day i.p. injections of FTY720 at a dose of 1 mg/kg, or of vehicle control containing 2.5% ethanol and 2% β-cyclodextrin. FTY720 was administered for a period of 11 days. Cells from spleen, liver and lung were analyzed one day after the final FTY720 injection.

### Multiplex gene expression analysis

γδ T cells were sorted from the lung and liver of TCRα^-/-^ mice (n=10), using the TCR γδ T cell Isolation kit (Miltenyi Biotec). RNA was extracted using the Nucleospin RNAII kit (Macherey Nagel), was quantified with a NanoDrop spectrophotometer (Promega), and was qualified with the 2100 Bioanalyzer System (Agilent). The nCounter GX analysis system (NanoString) was utilized according to the manufacturer’s directions to quantify RNA expression of 547 genes on the nCounter® Immunology Panel.

### CDR3 TCRδ (TRD) high-throughput sequencing

RNA was prepared from blood (250 µl) of individual mice using QIAmp RNA Blood kit (Qiagen) and NucleoSpin RNA blood kit (Macherey Nagel), or from sorted γδ T cells as above. cDNA was generated performing a template switch anchored RT-PCR. RNA was reverse transcribed via a template-switch cDNA reaction using 5’ CDS oligo (dT), a template-switch adaptor (5’-AAGCAGTGGTATCAACGCAGAGTACATrGrGrG) and the Superscript II RT enzyme (Invitrogen). The cDNA was then purified using AMPure XP Beads (Agencourt). Amplification of the TRD region was achieved using a specific TRDC primer (5’-*GTCTCGTGGGCTCGG*AGATGTGTATAAGAGACAGAAAACAGATGGTTTGGCCGGA, adapter in italic) and a primer complementary to the template-switch adapter (5’ *TCGTCGGCAGCGTCAGATGTGTATAAGAGACAG*AAGCAGTGGTATCAACGCAG, adapter in italic) with the KAPA Real-Time Library Amplification Kit (Kapa Biosystems). Adapters were required for subsequent sequencing reactions. After purification with AMPure XP beads, an index PCR with Illumina sequencing adapters was performed using the Nextera XT Index Kit. This second PCR product was again purified with AMPure XP beads. High-throughput sequencing of the generated amplicon products containing the TRG and TRD sequences was performed on an Illumina MiSeq platform using the V2 300 kit, with 150 base pairs (bp) at the 3′ end (read 2) and 150 bp at the 5′ end (read 1) [at the GIGA center, University of Liège, Belgium].

Raw sequencing reads from fastq files (read 1 and read 2) were aligned to reference V, D and J genes from GenBank database specifically for ‘TRD’ to build CDR3 sequences using the MiXCR software version 3.0.13 [70]. Default parameters were used except to assemble TRDD gene segment where 3 instead of 5 consecutive nucleotides were applied as assemble parameter. CDR3 sequences were then exported and analyzed using VDJtools software version 1.2.1 using default settings [71]. Sequences out of frame and containing stop codons were excluded from the analysis. Note that the nucleotype lengths generated by VDJtools include the C and V ends of the CDR3 clonotypes. The degree of TCR repertoire overlap between two different samples was analyzed using the overlap F metrics calculated with the software package VDJtools (https://vdjtools-doc.readthedocs.io/en/master/index.html)..

### Statistical analysis

Statistical studies were performed using GraphPad Prism 6 software and indicated in the Figure legends (*P <0,05; **P <0.01 ***P <0.001 and ****P <0.0001).

## AKNOWLEDGMENTS

We thank Isabelle Douchet at Immunoconcept for technical assistance during mice dissection.

## FIGURE CAPTIONS

**S1 Fig.**
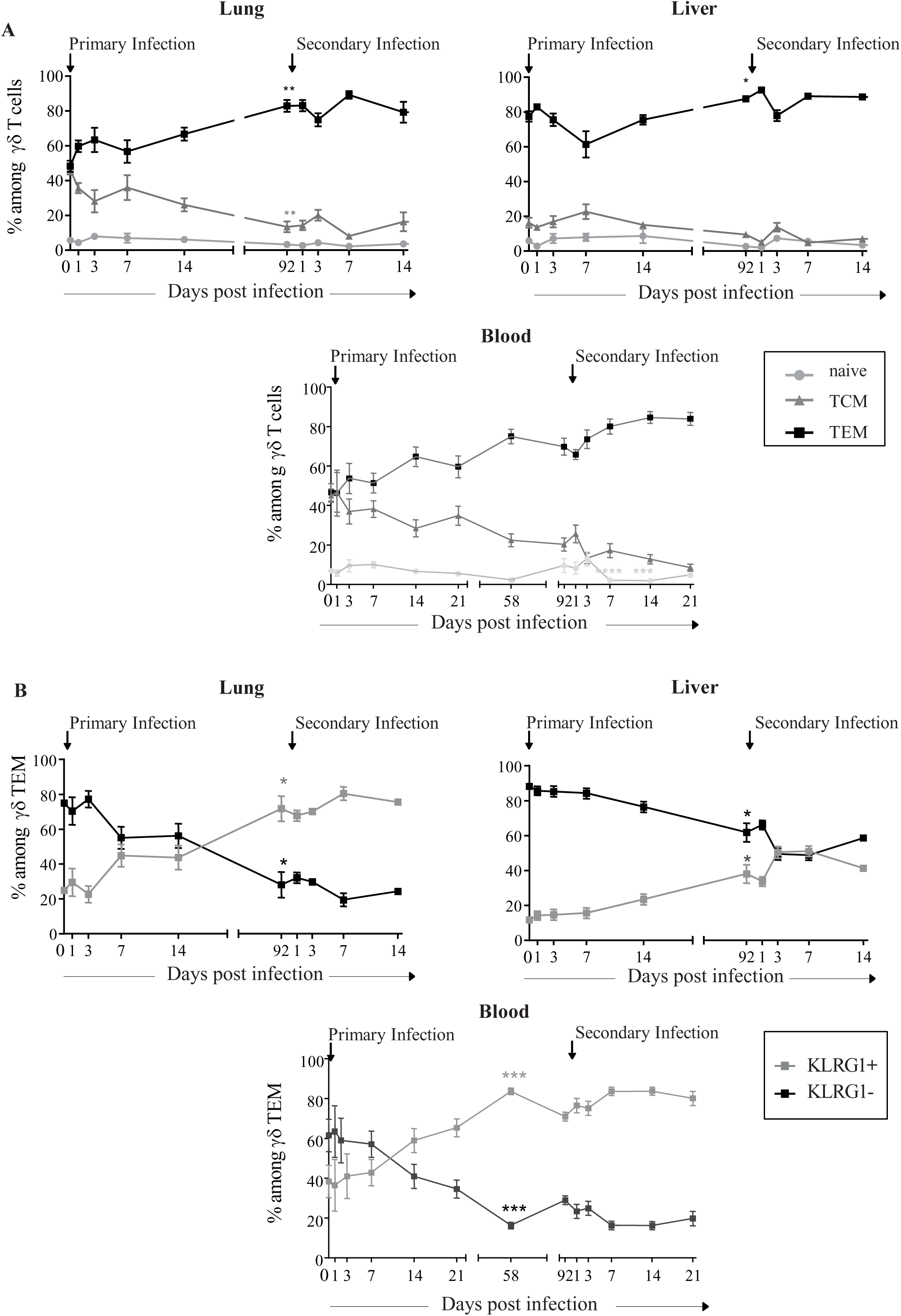
Kinetic evolution of the proportion of γδ T cell memory subsets in organs. TCRα-/- mice (n=25) were infected with MCMV (2.10^3^ PFU) at day 0, or left uninfected (n=25). 3 months later, uninfected mice were primarily infected, and infected mice were re-challenged with similar dose of MCMV. (A) Percentages of naïve (CD44-CD62L+), TCM (CD44+CD62L+) and TEM (CD44+CD62L-) cells among γδ T lymphocytes from organs. (B) Percentages of KLRG1+ and KLRG1-cells among γδ TEM. Data represent the mean percentages +/- SEM from 5 mice.

**S2 Fig.**
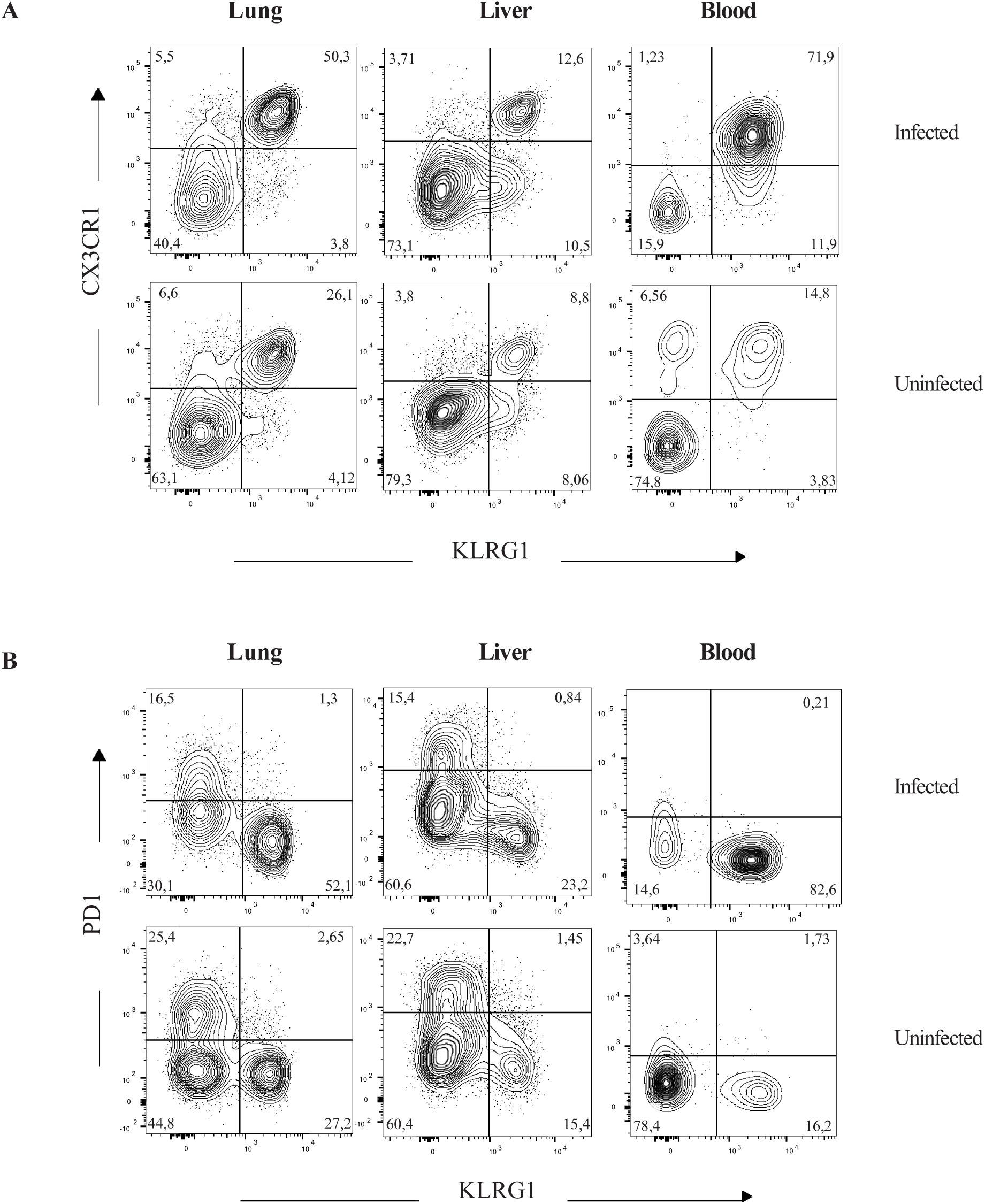
Phenotypic analysis of γδ TEM from d92 MCMV-infected and age- matched control mice. (A) Co-expression of CX3CR1 and KLRG1 on γδ TEM from organs and blood of d92 MCMV-infected and age-matched control mice. (B) Mutually exclusive expression of KLRG1 and PD1 γδ TEM from organs of d92 MCMV-infected and age-matched control mice. Numbers show percentages of each marker among γδ TEM. One representative mouse is shown for each (control and long-term MCMV infected) group of mice.

**S3 Fig.**
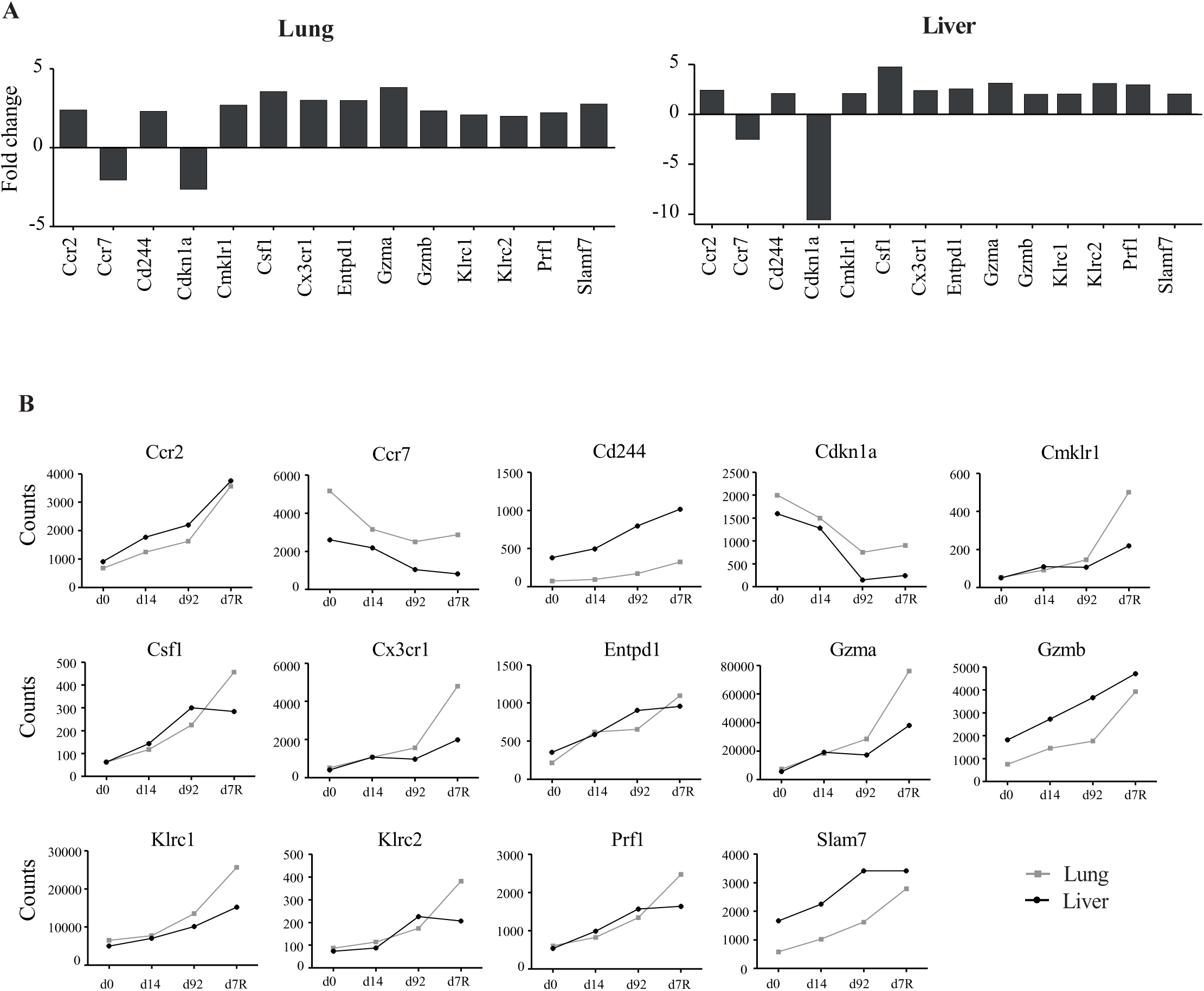
Multiplex gene expression analysis of immunology markers expressed by MCMV-induced γδ T cells. γδ T cells were sorted from lung and liver of TCRα-/- mice (n=10) at d0, d14 or d92 post MCMV infection, and day 7 post-reinfection (d7R). Analyses were performed with nSolver™ Analysis Software (NanoString). (A) Histograms represent transcripts shared by liver and lung, and whose d92/d0 ratios were >2 or < -2 (minimum counts=100). (B) Counts evolution of theses transcripts at day 0, 14, 92 post-infection (d0, d14, d92), and day 7 post reinfection (d7R).

**S4 Fig.**
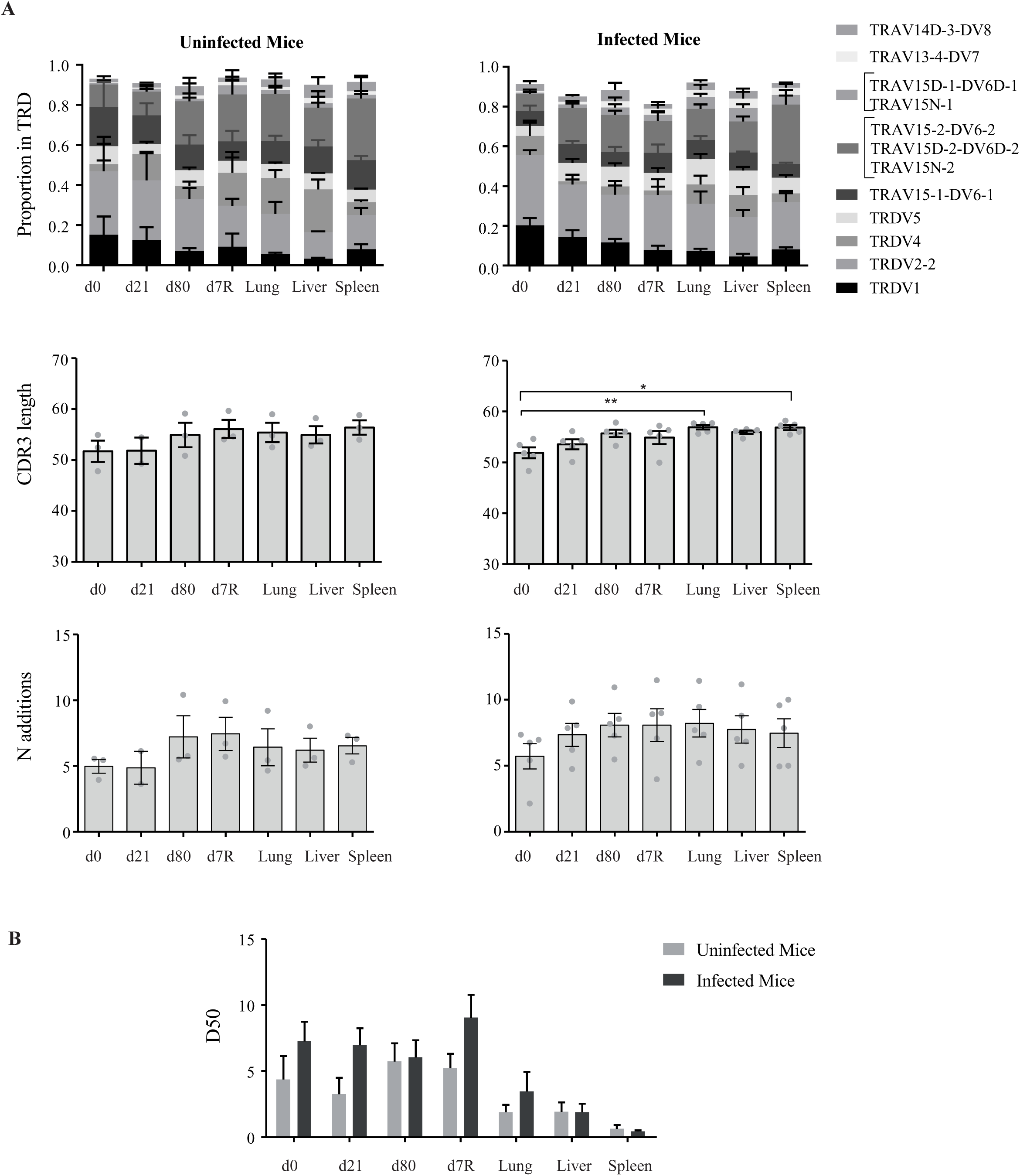
CDR3δ repertoire analysis. TCRα-/- mice (n=5) were infected with MCMV. Age matched uninfected TCRα-/- mice (n=3) were used as controls. Blood was drawn at different time intervals post infection. At day 7 post reinfection, blood and organs were analyzed. (A) Comparison between infected (right panels) and age-matched control mice (left panels), of the CDR3 TRD repertoire of blood samples at d0, 21, 80 post-infection (d0, d21, d80) and at d7 post-reinfection (d7R), and from organs at d7R. (Upper panels) TRDV usage distribution, (Middle panels) CDR3 length in nucleotides (including the codons for C-start and F-end residues), each dot represents the weighted mean of an individual sample. (Lower panels) Number of N additions, each dot represents the weighted mean of an individual sample. (B) Percentage of unique clonotypes required to account for 50 % of the total repertoire in infected or age-matched control mice. Statistical test was 1-way ANOVA.

**S5 Fig.**
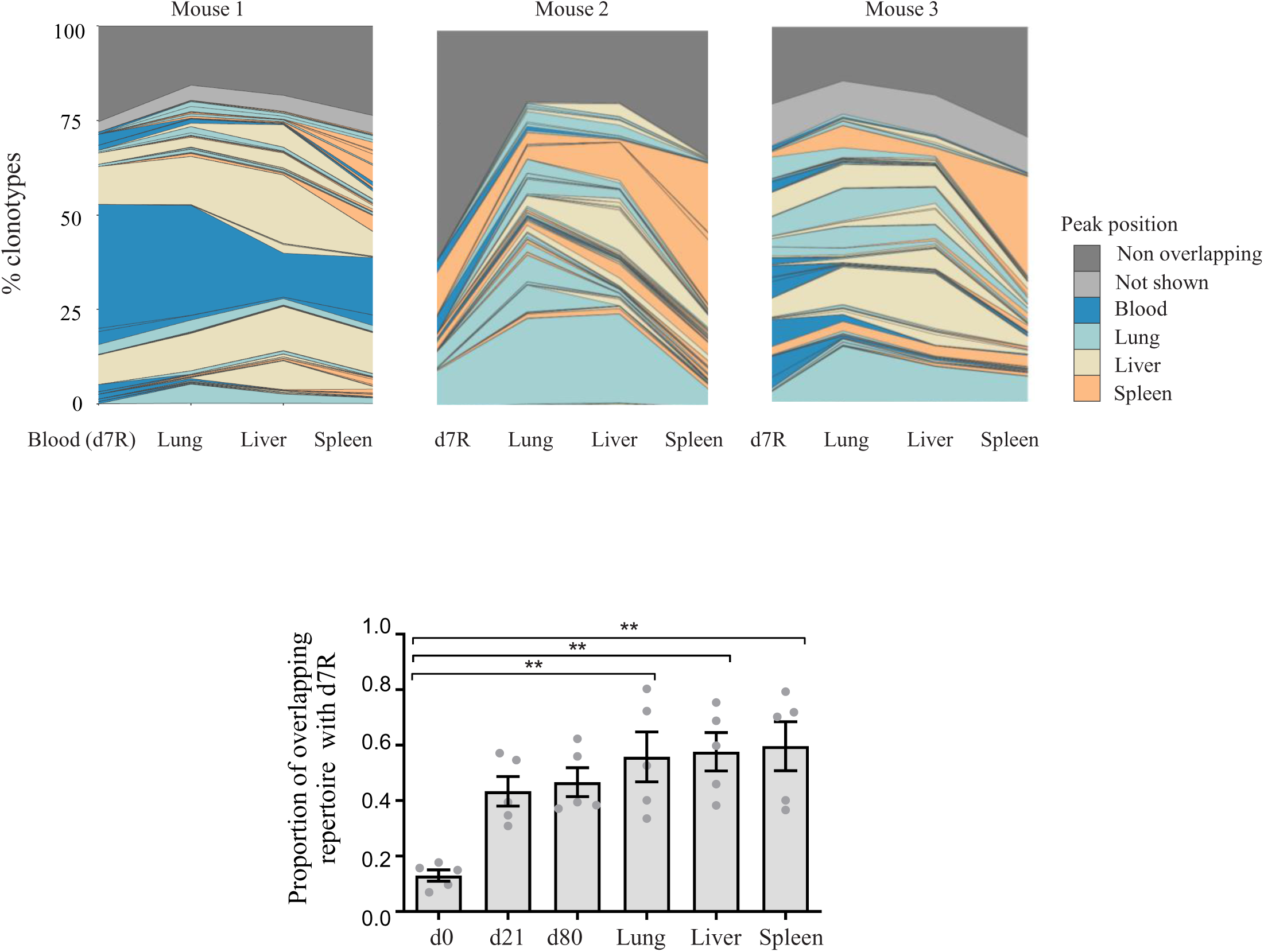
Tracking of shared clonotypes after reinfection in blood and organs. TCRα-/- mice (n=5) were infected with MCMV and reinfected 3 months later. At day 7 post reinfection (d7R), blood and organs were analyzed. (Upper panels) Examples of clonotype tracking stackplots for 3 infected mice: detailed profiles for top 100 clonotypes, as well as collapsed (“NotShown” in dark gray) and non-overlapping (light gray) clonotypes found in blood and organs at d7R. Clonotypes are colored by the peak position of their abundance profile. The colors are matched for each mouse samples but not between the different mice. (Lower panels) Overlap frequencies of the D7R repertoire compared to the indicated column. Each dot corresponds to one pair comparison and one mouse. Statistical test was 1-way ANOVA.

**S6 Fig.**
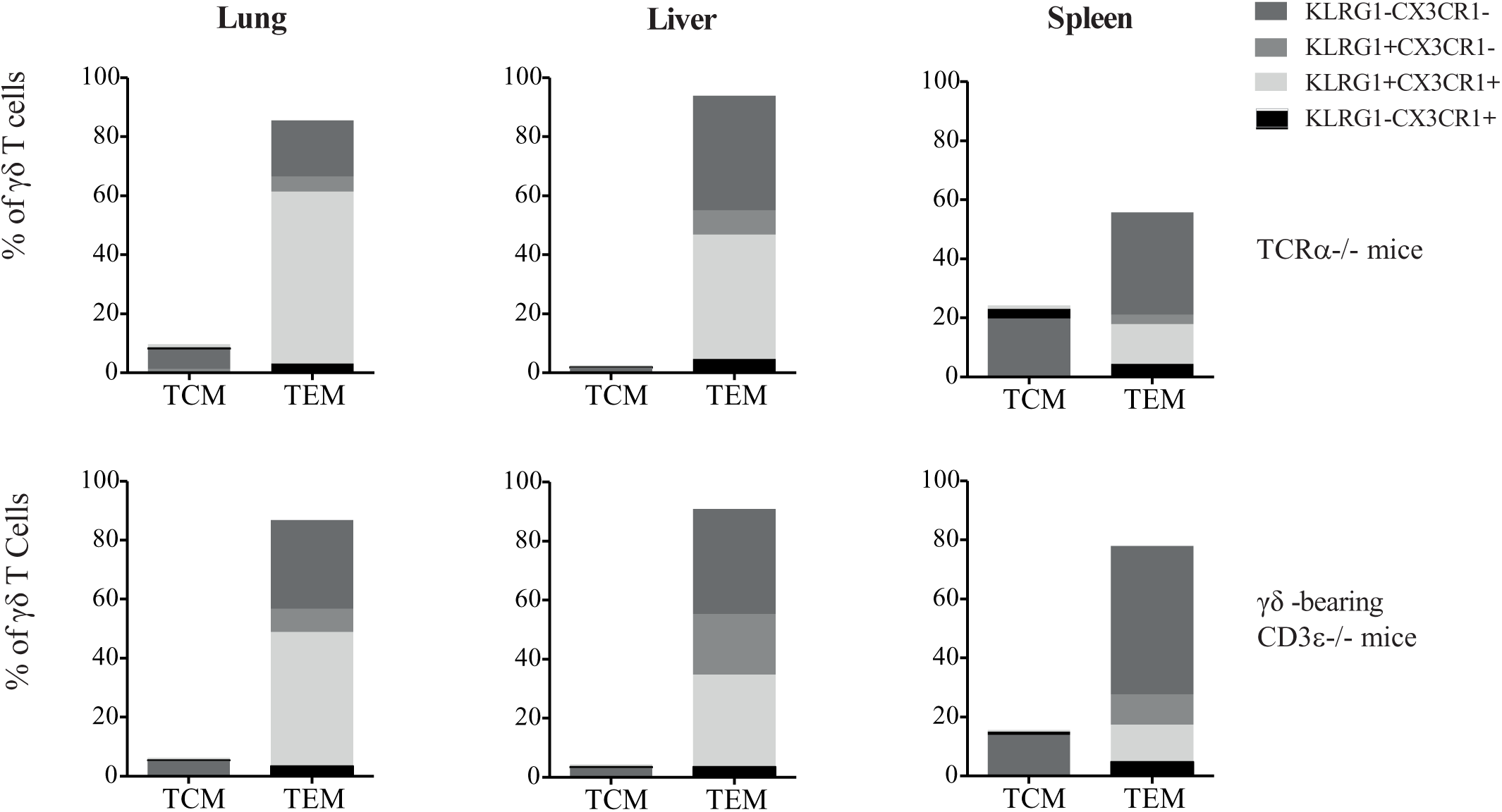
Repartition of KLRG1/CX3CR1 expressing γδ T cell subtypes in γδ-bearing CD3ɛ-/- mice infected with MCMV and in d92-infected TCRα-/- mice. Long-term MCMV-induced γδ T cells were sorted from the spleen of TCRα-/- mice and transferred into CD3ɛ-/- mice that were subsequently infected with MCMV. (Upper panels) Percentages of γδ TCM and TEM were determined in organs from 5 d92-infected TCRα-/- mice. (Lower panels) Percentages of γδ TCM and TEM were determined in organs from 8, γδ bearing CD3ɛ-/- mice that had survived until d130. The proportion of indicated subtypes within total TCM and TEM is shown by shades of grey.

**S7 Fig.**
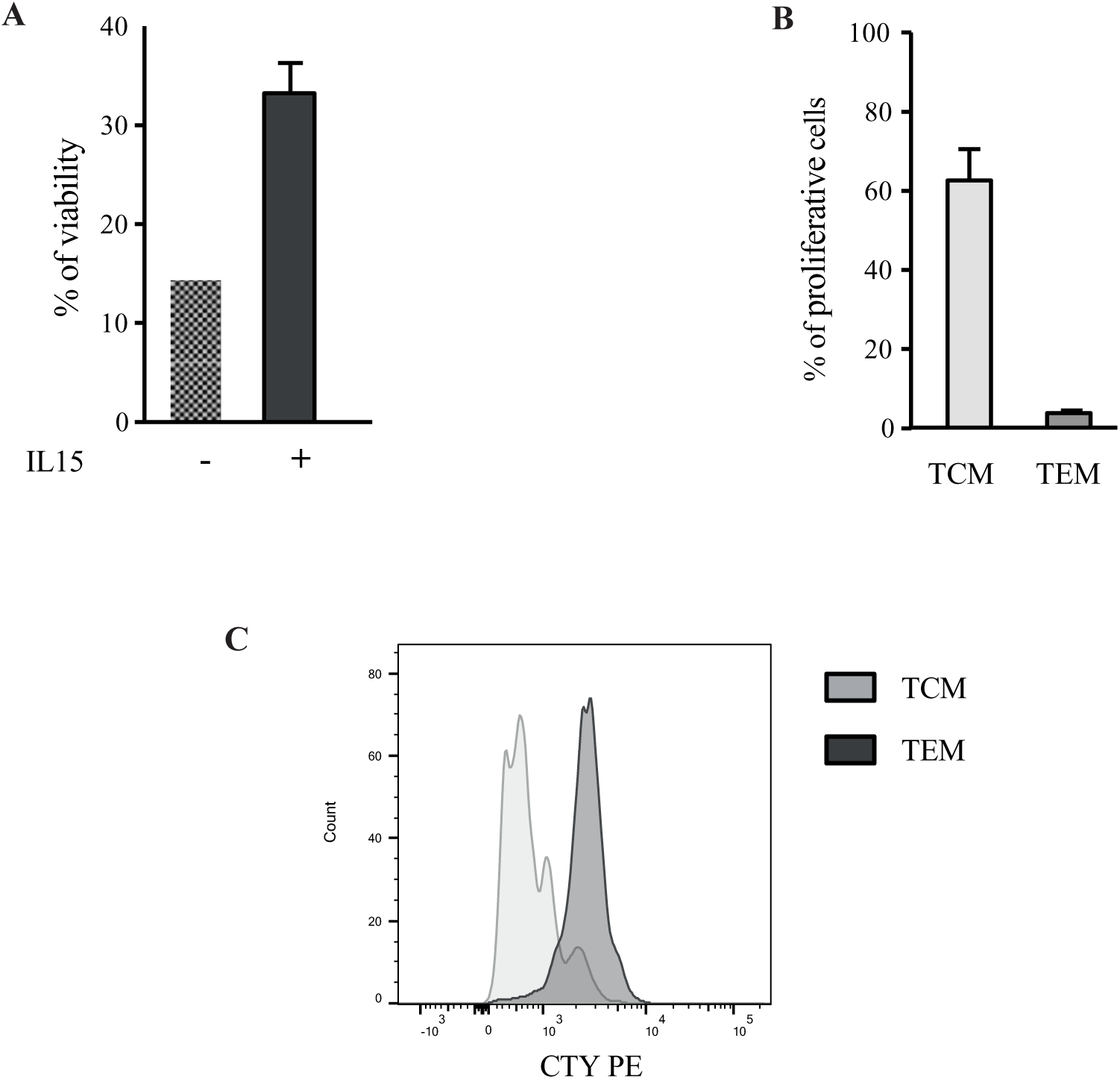
γδ TCM are more proliferative than γδ TEM in response to IL15. Total γδ T cells were sorted from the spleen of 3 months infected TCRα-/- mice. Cells were labelled with CTY and cultured with or without IL15 (200 ng/µl) for 3 days. (A) Percentages of viable cells among γδ T cells cultured in the absence or presence of IL15. (B) Percentages of proliferative γδ TCM or TEM after 3 days of culture with IL15 (C) Representative histograms of cellular proliferation for TCM (light grey) and TEM (dark grey) from d92-infected TCRα-/- mice. Data are representative of 3 independent experiments.

## Notes

### Competing Interest Statement

The authors have declared no competing interest.

